# Behavior-coupled single-cell transcriptomics reveals trafficking protein-dependent regulation of tumor cell migration

**DOI:** 10.64898/2026.10.15.759353

**Authors:** Reza Ghadiri Rad, Junfeng Huang, Maria Teresa Lopez-Cascales, Li You, Joleen J.H. Traets, Domenico Bellomo, Ting-Chun Chou, Wesley Bennik, Jasmin Singh, Yasin Begce, Jose Angelito U Hardillo, Disha Vadgama, Thierry van den Bosch, Vera van Dis, Miao-Ping Chien

**Affiliations:** Department of Molecular Genetics, Erasmus University Medical Center, Rotterdam, The Netherlands.; Erasmus MC Cancer Institute, The Netherlands; Oncode Institute, Utrecht, The Netherlands; SkylineDx B.V., Rotterdam, The Netherlands; Department of Otorhinolaryngology, Head and Neck Surgery, Erasmus University Medical Center, Rotterdam, The Netherlands; Department of Pulmonary Medicine, Erasmus University Medical Center, Rotterdam, The Netherlands; Department of Pathology and Clinical Bioinformatics, Erasmus University Medical Center, Rotterdam, The Netherlands

**Author notes:** Senior author. These authors contributed equally (Reza Ghadiri Rad, Junfeng Huang).

## Abstract

Aggressive cellular migration underlies cancer invasion and metastatic dissemination, yet the molecular programs defining highly migratory tumor cell states remain poorly characterized. In head and neck squamous cell carcinoma (HNSCC), metastatic progression is associated with poor clinical outcome, motivating method developments that link migratory behavior to underlying mechanisms. Here, we present behavior-coupled single-cell sequencing (BC-seq), a live-cell imaging-guided transcriptomic platform that enables real-time identification, isolation, and profiling of individual cells based on migratory behavior.

Applying BC-seq to 10 HPV-negative oral squamous cell carcinoma (OSCC), the most common HNSCC subtype, we distinguished fast-migrating cells (FMCs) from slow-migrating cells (SMCs) and identified transcriptional programs associated with aggressive migration. Mechanistically, BC-seq uncovered cholesterol-regulated signaling pathways and associated trafficking proteins involving SREBP2, STARD4, Cavin1/3, and VEGFR2 that promotes actin cytoskeletal remodeling and cellular motility. Disruption of STARD4-mediated cholesterol trafficking or Cavin1/3-dependent VEGFR2 nuclear translocation significantly impaired cell migration, linking cholesterol metabolism to migratory machinery.

We next examined whether molecular programs associated with aggressive migration are reflected in clinical outcomes. A 15-gene expression signature derived from FMCs predicted distant metastasis with 83.6% accuracy in an independent OSCC cohort, outperforming conventional clinicopathological risk stratification. Together, our study establishes BC-seq as an approach for integrating dynamic cellular behavior with single-cell transcriptomics to uncover mechanisms of aggressive migration and clinically relevant molecular signatures.

## Introduction

Cellular migration is a fundamental biological process that underlies tissue development, wound repair, and immune surveillance, and its dysregulation is a hallmark of cancer invasion and metastatic dissemination^1^. In solid tumors, the ability of cancer cells to adopt highly migratory phenotypes enables local invasion, intravasation, and eventual colonization of distant organs^2^. In head and neck squamous cell carcinoma (HNSCC), aggressive migratory behavior is closely associated with disease progression and poor clinical outcome, particularly in cases that develop distant metastases^3^. At the cellular level, migratory capacity is governed by coordinated regulation of cytoskeletal dynamics, membrane trafficking, and signaling pathways that respond to microenvironmental cues^4^. Increasing evidence suggests that tumor cells do not migrate uniformly; instead, highly migratory behavior is often confined to rare subpopulations that transiently acquire enhanced motility^5^. Capturing these aggressive migratory states and linking them to underlying molecular programs remains a central challenge in metastasis research.

Single-cell RNA sequencing (scRNA-seq) has transformed the study of tumor heterogeneity by enabling transcriptional profiling at cellular resolution^6,7^. Across multiple cancer types, scRNA-seq has uncovered rare malignant cell states and delineated tumor-stromal interactions associated with invasion and metastatic dissemination. In HNSCC, Puram et al. identified malignant and stromal subpopulations linked to invasive behavior^8^, while analogous studies in colorectal and breast cancers described stem-like and invasion-associated states enriched in metastatic lesions^9,10^. However, most single-cell studies rely on static transcriptional snapshots and therefore provide limited insight into dynamic cellular behaviors such as migration.

Among the transcriptional programs associated with invasive behavior, epithelial-to-mesenchymal transition (EMT) and partial EMT (p-EMT) states have been repeatedly implicated in cancer progression^8,11^. EMT-related signatures have also been detected in circulating tumor cells from patients with metastatic disease ^12,13^, and p-EMT programs correlate with invasion and nodal involvement in HNSCC^8^. At the same time, EMT-associated programs appear to be highly dynamic and context dependent, with variable persistence across tumors and disease stages^14^. These observations suggest that aggressive migratory behavior may not always align with stable transcriptional identities, highlighting the need for approaches that directly link functional migration phenotypes to molecular states.

Motivated by this, we hypothesized that aggressive migratory capacity may be concentrated within rare, transient tumor cell states that are best defined by behavior. Conceptual models and experimental studies have proposed that such dynamic phenotypic states can contribute disproportionately to metastatic dissemination^15,16^, but systematically capturing and molecularly characterizing these cells has remained technically challenging. To address this limitation, we developed BC-seq (behavior-coupled single-cell sequencing), a live-cell imaging-guided transcriptomic platform that enables real-time identification, isolation, and profiling of individual cells based on migratory behavior. BC-seq directly links dynamic cellular phenotypes to transcriptional programs, allowing interrogation of the molecular mechanisms underlying aggressive migration. Using this approach, we distinguished fast-migrating cells (FMCs) from slow-migrating cells (SMCs) in HPV-negative oral cavity HNSCC, also known as oral squamous cell carcinoma (OSCC), the most common HNSCC subtype.

Mechanistically, BC-seq revealed a cholesterol-regulated signaling cascade and associated trafficking proteins involving SREBP2, STARD4, Cavin1/3, and VEGFR2 that orchestrates actin cytoskeletal remodeling and cellular migration. Functional perturbation of STARD4-mediated cholesterol trafficking and Cavin1/3-dependent VEGFR2 nuclear translocation significantly impaired cell migration, establishing a direct mechanistic link between cholesterol metabolism and migratory machinery. These findings identify a previously uncharacterized cholesterol-VEGFR2 axis regulating aggressive migration in OSCC and suggest broader relevance to tumor cell motility.

Given the close association between migratory behavior and metastatic dissemination, we next asked whether molecular programs defining aggressively migrating cells could be linked to clinical outcomes. From FMCs, we derived a 15-gene expression signature that predicts distant metastasis with 83.6% accuracy in an independent cohort of 81 OSCC patients, outperforming conventional clinicopathological risk stratification approaches, which typically achieve 20-30% sensitivity^17,18^.

Together, our study introduces a behavior-guided framework for dissecting tumor heterogeneity, integrating dynamic cellular phenotypes with single-cell transcriptomics to uncover mechanistic insights of aggressive migration and clinically informative biomarkers of metastatic risk.

### Behavior-coupled single-cell transcriptomics

Phenotypic plasticity enables cancer cells to adopt a range of functional states that contribute to tumor progression, therapeutic resistance, and metastatic dissemination^19^. In particular, aggressive migratory behavior often arises transiently within small subsets of tumor cells. However, capturing such migratory phenotypes in a behavior-resolved and cell type-specific manner remains technically challenging, as they are heterogeneous, transient, and difficult to identify using static molecular profiling approaches. To overcome these limitations, we developed BC-seq to enable real-time identification, isolation, and molecular characterization of individual OSCC cells based on migratory behavior.

BC-seq begins by imaging tens of thousands of single cells pre-incubated with photoactivatable dyes using a high-content single-cell screening microscope (Fig. 1a, b; Supplementary Fig. S1; Methods). To minimize patient-specific heterogeneity and identify shared molecular mechanisms, BC-seq was performed on 10 patient-derived samples. Real-time image analysis is then performed to track individual cell trajectories and identify subpopulations exhibiting fast or slow migratory behavior (Fig. 1a, b; Methods). Once identified, target cells are selectively phototagged using a 405 nm laser directed by galvo mirrors integrated into the microscope (Fig. 1a, b; Methods). These behaviorally defined cells are then isolated via fluorescence-activated cell sorting (FACS) and subjected to single-cell RNA sequencing, enabling precise molecular profiling of transient cellular phenotypes.

**Figure 1.**
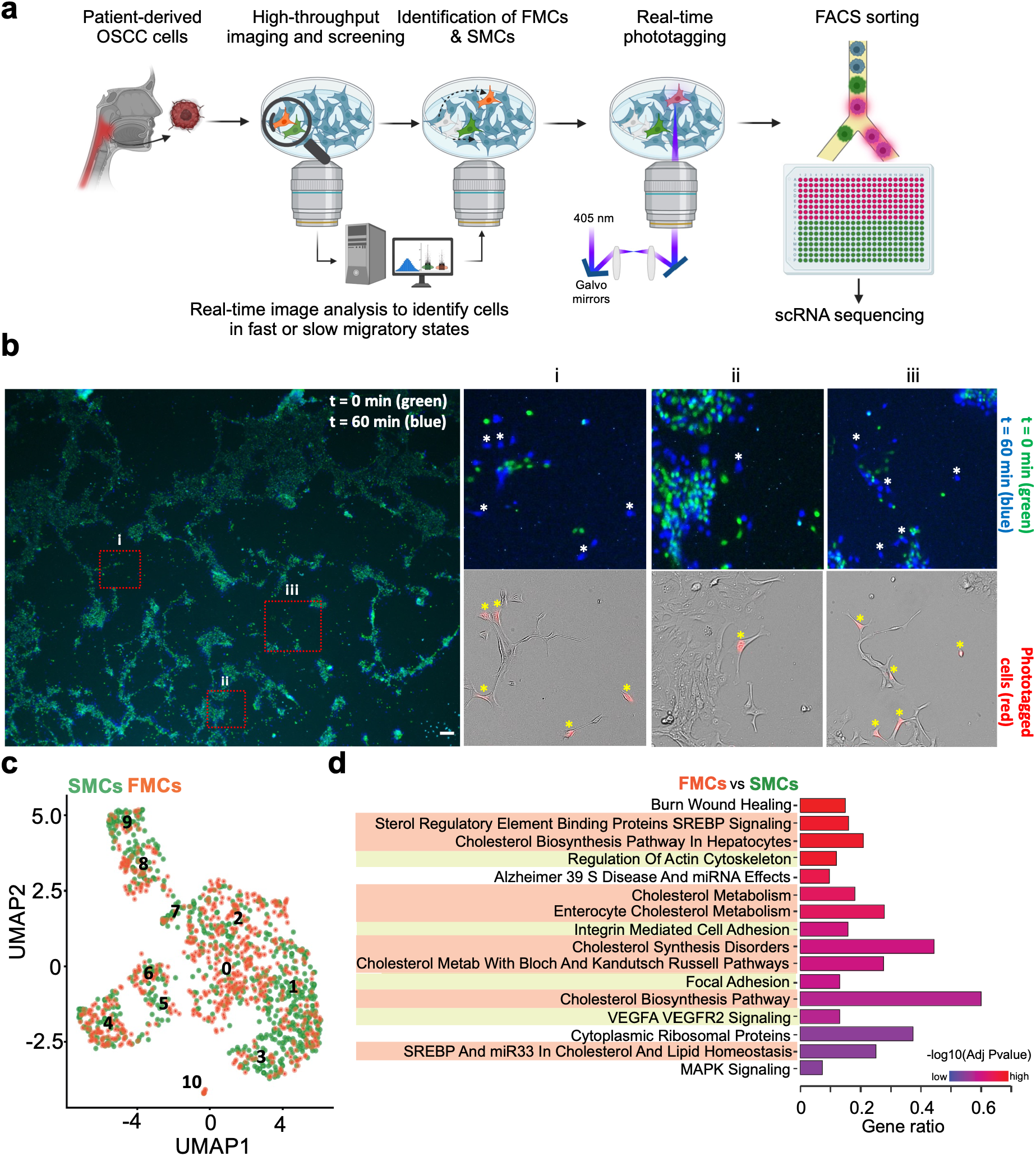
BC-seq profiling of fast- and slow-migrating OSCC cells. **a.** Schematic overview of the BC-seq pipeline. Patient-derived primary OSCC cultures are imaged using a custom-built high-throughput screening microscope. Real-time image analysis identifies cells of interest (e.g., FMCs – fast migrating cells) which are then photolabeled, isolated by FACS, and subjected to single-cell sequencing and profiling. **b**. Overlay of OSCC cell positions at t=0 (green) and t=60 minutes (blue). Insets i-iii: Eleven representative FMCs (indicated by asterisks) identified through image analysis and labelled via phototagging. Scale bar, 200 μm. **c**. A UMAP plot illustrates cells in either fast-or slow-migrating states (FMCs or SMCs). The numbers indicate the 11 clusters (Clusters 0-10) identified by Seurat. **d**. WikiPathways analysis of differential genes from total FMCs cells versus SMCs control cells. Fisher’s exact test was applied for the Enrichr pathway enrichment analysis.

BC-seq enables the enrichment of selected cell types of interest, in this case FMCs and SMCs, and allows for the direct linking of fast or slow cell migration to transcriptomic profiles without interference from non-specific cells. As a result, cell type-or phenotype-specific gene profiles are greatly enhanced by BC-seq, thereby facilitating the identification of relevant molecular mechanisms. The ability to annotate phenotypes in advance is particularly critical when distinct phenotypic subpopulations share predominant transcriptomic profiles. In such cases, traditional clustering analysis often fails to distinguish the desired phenotypes (Fig. 1c). This behavior-informed profiling revealed a distinct enrichment of cholesterol synthesis and metabolism pathways in FMCs compared to SMCs (Fig. 1d-highlighted in orange). Notably, the cholesterol synthesis and metabolism signaling pathways were absent in the expression profiles of any cell subgroups identified through non-BC-seq-mediated scRNA-seq and clustering analysis (Supplementary information, Fig. S2). This finding underscores the importance of using phenotypically anchored approaches to uncover state-specific gene programs.

### SREBP2-mediated cholesterol synthesis signaling is correlated with fast cell migration

Metabolic reprogramming is essential for malignant transformation and tumor progression, driving processes like invasion and metastasis^20–23^. To confirm the role of cholesterol synthesis and metabolism signaling in promoting fast cellular migration in OSCC, we monitored migratory behavior under cholesterol-depleted conditions using methyl-β-cyclodextrin (MβCD) (Fig. 2a, top) and under cholesterol-enriched conditions (Fig. 2a, bottom). We observed a significant reduction in migration under cholesterol-depleted conditions and a corresponding increase under cholesterol-enriched conditions, confirming that cholesterol signaling contributes to enhanced cellular migration.

**Figure 2.**
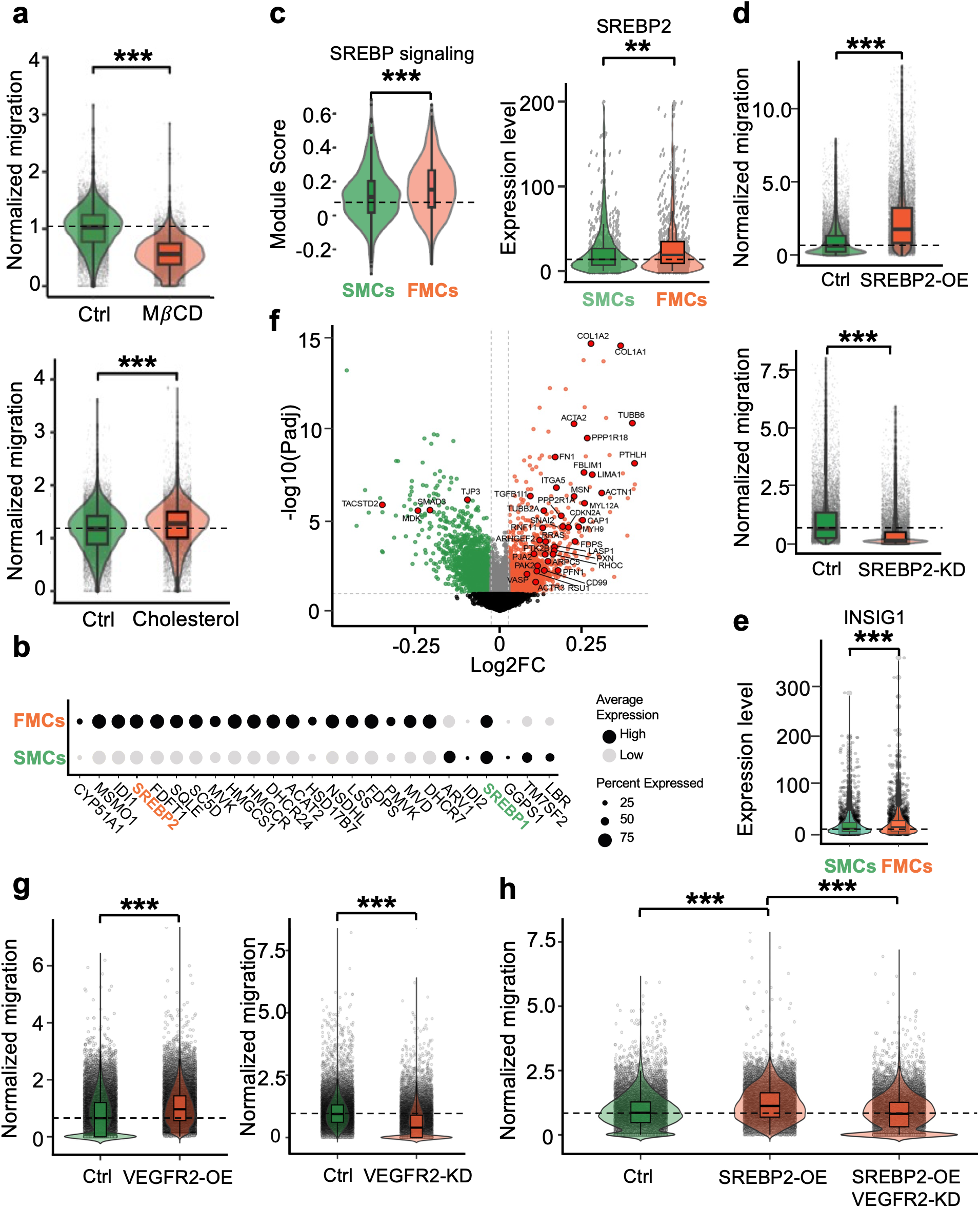
SREBP2-driven cholesterol synthesis and VEGFR2-mediated signaling promote cell migration. **a.** (Top panel) A violin plot displays normalized migration of cells under the cholesterol-depleted condition (MβCD, right) compared to the control (left). (Bottom panel) A violin plot displays the normalized migration of cells under the cholesterol-enriched condition (Cholesterol, right) compared to the control (left); MβCD, methyl-β-cyclodextrin; Cholesterol, cholesterol-enriched condition. **b.** Gene expression levels of cholesterol metabolism in both FMCs versus SMCs are displayed in a bubble plot. **c.** (Left panel) A violin plot shows a module score of SREBP signaling for SMCs and FMCs. (Right panel) A violin plot shows gene expression level of SREBP2 for SMCs and FMCs. **d.** (Top panel) A violin plot shows normalized migration of cells under SREBP2-overexpression (OE). (Bottom panel) A violin plot shows normalized migration of cells under SREBP2-Knock down (KD). **e.** A violin plot shows gene expression level of INSIG1. **f.** A volcano plot shows DEGs between FMCs and SMCs, with highlighted genes associated with actin cytoskeleton remodeling and focal adhesion. **g.** (Left panel) A violin plot shows the normalized migration of cells under control (Ctrl) and VEGFR2-overexpression (OE) conditions. (Right panel) A violin plot shows the normalized migration of cells under control (Ctrl) and VEGFR2-knockdown (KD) conditions. **h.** A violin plot shows the normalized migration of cells under control (Ctrl), SREBP2-overexpression (OE), and combined SBREP2-OE with VEGFR2-KD conditions. For a, c, d, e, g and h, *P* values were calculated using the Wilcoxon rank sum test with Bonferroni correction.

Among the upregulated genes associated with cholesterol synthesis and metabolism, many are intricately linked to the sterol regulatory element-binding protein (SREBP) signaling pathway (Fig. 1d, 2b, c). SREBP transcription factors play a pivotal role in regulating both lipid and cholesterol metabolism^24^. Specifically, SREBP2 is the main regulator of cholesterol biosynthesis and homeostasis^24^, while SREBP1 primarily governs lipid synthesis. Unlike SREBP1 (Supplementary information, Fig. S3), SREBP2 was uniquely upregulated in FMCs (Fig. 2b, c). To confirm the role of SREBP2-mediated cholesterol synthesis and metabolism signaling in driving fast migration, we conducted both SREBP2-overexpression (OE) and SREBP2-knockdown (KD) assays (Fig. 2d; The qPCR results of SREBP2 in the SREBP2-OE and SREBP2-KD assays are shown in Supplementary information, Fig. S4). These experiments revealed increased migratory speed under SREBP2-OE conditions and reduced migration under SREBP2-KD conditions, underscoring the critical role of SREBP2 in driving fast cell migration (Fig. 2d). Furthermore, another cholesterol synthesis and metabolism-related gene, INSIG1, was found to be upregulated in FMCs (Fig. 2e). INSIG1 is a known negative regulator of cholesterol biosynthesis, with high cholesterol production often resulting in increased INSIG1 expression^25^.

### Cavin 1 and 3-mediated caveolae formation enhances VEGFR2 nuclear translocation and activates serum response factor-mediated cell migration

In addition to the SREBP2 signaling and cholesterol synthesis pathways, VEGFA-VEGFR2 and actin cytoskeleton-related signaling pathways were notably upregulated in FMCs, as revealed by pathway analysis (Fig. 1d; highlighted in yellow). The gene program related to actin cytoskeleton remodeling and focal adhesion is clearly upregulated in FMCs, including genes such as COL1A1, COL1A2, ACTA2, FN1, ACTN1, TGFB1I1, FBLIM1, VASP, and PPP1R18 (Fig. 2f). VEGFA-VEGFR2 signaling has been shown to be involved in cell migration in both endothelial and epithelial cancer cells, affecting tumor functions independently of angiogenesis^26–28^. However, the mechanisms by which VEGFA-VEGFR2 signaling drives cell migration, and its link to SREBP2-mediated cholesterol synthesis, remain unclear.

Before addressing these questions, we first confirmed the role of VEGFR2-mediated cell migration in OSCC, we conducted both VEGFR2-overexpression (OE) and VEGFR2-knockdown (KD) assays (Fig. 2g; The qPCR results of VEGFR2 in the VEGFR2-OE and VEGFR2-KD assays are shown in Supplementary information, Fig. S5). A clear increase in migratory speed was observed under VEGFR2-OE conditions, while a reduction was seen under VEGFR2-KD conditions, confirming the role of VEGFR2 in driving cellular migration (Fig. 2g).

As VEGFR2 (vascular endothelial growth factor receptor 2), a member of the transmembrane receptor tyrosine kinase (RTK) family, is known to be involved in intracellular signaling cascades involved in cell motility and migration^26–28^, we hypothesized that SREBP2 acts as an upstream regulator of VEGFR2-mediated cell migration. We confirmed this relationship by assessing cellular migration in SREBP2-OE and SREBP2-OE combined with VEGFR2-KD conditions (Fig. 2h). Increased migration was observed in the SREBP2-OE condition, while a reversal of migration was seen when VEGFR2 was knocked down in the SREBP2-OE background (Fig. 2h).

To confirm that SREBP2-OE also leads to VEGFR2 protein upregulation, we performed immunofluorescence staining of VEGFR2 under SREBP2-OE and control conditions (Fig. 3a). As expected, VEGFR2 expression was clearly upregulated in the SREBP2-OE condition, but interestingly, it was unexpectedly localized to the nucleus. To determine whether this nuclear translocation was related to SREBP2-mediated signaling, we also performed VEGFR2 immunofluorescence staining under VEGFR2-OE alone conditions. Notably, nuclear translocation and expression intensity of VEGFR2 were significantly lower in this setting (Supplementary Fig. S6), confirming that VEGFR2 nuclear translocation is a downstream consequence of SREBP2 overexpression.

**Figure 3.**
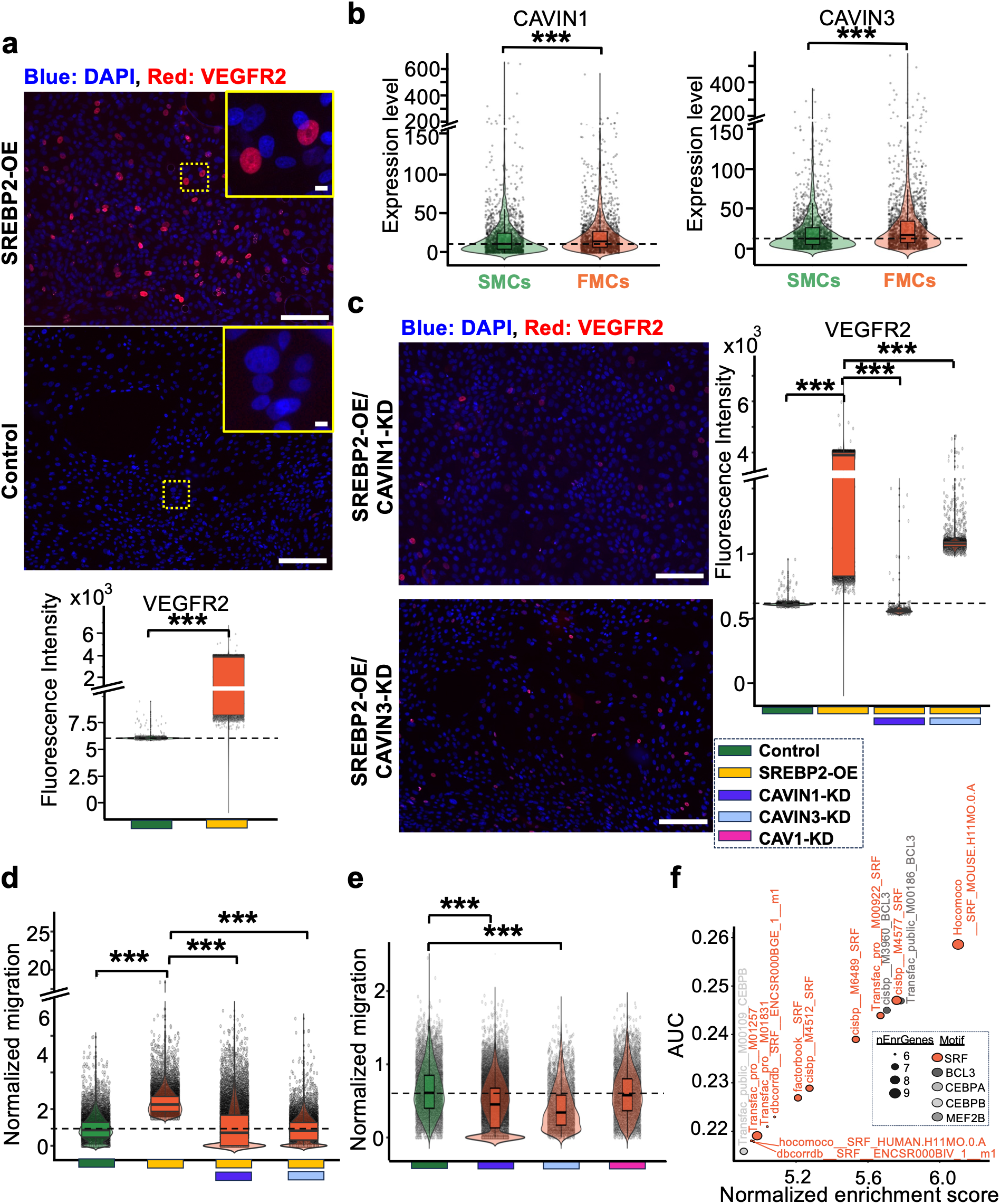
Cavin 1 and 3-mediated caveolae formation enhances VEGFR2 nuclear translocation and activates serum response factor-mediated cell migration in OSCC. **a.** (Top panel) Representative immunofluorescence images of OSCC cells stained with DAPI (blue) and VEGFR2 (red) under control and SREBP2-overexpression (OE) conditions. (Bottom panel) Fluorescence intensity of nuclear VEGFR2 under control (green) and SREBP2-OE (yellow) conditions. Scale bars = 50 μm; 5 μm for insets. b. A violin plot shows gene expression level of CAVIN1 (left) and CAVIN3 (right) for SMCs and FMCs. c. Representative immunofluorescence images of OSCC cells stained with DAPI (blue) and VEGFR2 (red) under combined SREBP2-OE with CAVIN1-knockdown (KD) conditions (top panel) or combined SREBP2-OE with CAVIN3-KD conditions (bottom panel). Scale bars = 50 μm. **d.** A violin plot shows normalized migration of cells under control (green), SREBP2-OE (yellow), combined SREBP2-OE with CAVIN1-KD (yellow & blue), or combined SREBP2-OE with CAVIN3-KD (yellow & cyan) conditions. **e.** A violin plot shows normalized migration of cells under control (green), CAVIN1-KD (blue), CAVIN3-KD (cyan), or CAV1-KD (pink) conditions. **f.** Motif enrichment analysis highlights serum response factor (SRF) as the most strongly predicted transcription factor (TF), shown in red. For a, b, c, d, e, *P* values were calculated using the Wilcoxon rank sum test with Bonferroni correction.

Nuclear translocation of other RTK family members, such as epidermal growth factor receptor (EGFR), has been previously reported and shown to regulate various cellular processes, including cell cycle progression and proliferation^29^. However, nuclear translocation of VEGFR2 remains poorly understood. We hypothesize that endocytic pathways, such as caveolae-mediated endocytosis^30^, which involve protein trafficking from the plasma membrane to the nuclear envelope, may facilitate and mediate VEGFR2’s nuclear translocation.

Among the genes linked to the caveolae signaling pathways, *CAVIN1* and *CAVIN3*, which encode caveolae associated protein 1 and 3 (Cavin 1 and 3), were uniquely upregulated in FMCs (Fig. 3b). We hypothesized that Cavin1 and Cavin3 are key transporters responsible for VEGFR2 nuclear translocation. To test this, we performed immunostaining of VEGFR2 under SREBP2-OE conditions combined with CAVIN1 or CAVIN3 knockdown (KD). Notably, VEGFR2 nuclear translocation was clearly abolished upon CAVIN1 or CAVIN3 knockdown (Fig. 3c). To further confirm that Cavin1 and Cavin3 not only facilitate VEGFR2 nuclear translocation but also play a functional role in facilitating fast cellular migration, we assessed migratory behavior under SREBP2-OE alone and SREBP2-OE with CAVIN1/CAVIN3-KD conditions (Fig. 3d). A significant reduction in migratory speed was observed under combined SREBP2-OE with CAVIN1-KD or CAVIN3-KD, confirming that Cavin1/3-dependent migration occurs downstream of SREBP2 upregulation. These data also suggest a connection between upregulated cholesterol levels and the formation of caveolae microdomains, aligning with recent studies showing that membrane cholesterol is critical for maintaining membrane fluidity and facilitating the formation of membrane protrusions and caveolae microdomains necessary for intracellular signaling^31^.

To further investigate whether caveolae-mediated fast migration is specifically driven by Cavin 1 and 3, as opposed to other caveolae-associated proteins like Caveolin 1 (not found in our differential profiles), we conducted individual genetic knockdowns of CAVIN1, CAVIN3 and CAV1 (CAV1 encodes Caveolin 1) (Fig. 3e). Strikingly, the knockdown of CAVIN 1 and 3 resulted in a pronounced reduction in cell migration, whereas CAV1 knockdown caused only a modest decrease (Fig. 3e; The qPCR results for CAVIN 1 and 3 and CAV1 gene expression after KD are shown in Supplementary information, Fig. S7). These findings highlight the critical role of Cavin 1 and 3, known for regulating caveolar trafficking^32^, in facilitating cholesterol upregulation-mediated migration.

Building on the above findings, we hypothesized that increased cholesterol production, mediated by SREBP2, in OSCC cells in a fast migratory state, promotes cholesterol incorporation into the plasma membrane, thereby facilitating the formation of caveolae membrane microdomains. This process, in turn, enhances Cavin1/3-mediated endocytosis, leading to nuclear translocation of VEGFR2 and activation of VEGFA-VEGFR2-mediated cell migration.

To understand the mechanisms by which VEGFA-VEGFR2 signaling drives cell migration, specifically how VEGFR2 nuclear translocation contributes to this process, we first identified potential transcription factors (TFs) involved. Nuclear translocation of RTKs, especially EGFR, often induces activation of TFs by acting as a transcriptional co-activator^29^. To identify potential TFs activated by nuclear VEGFR2, we performed transcription factor prediction using RcisTarget^33^ (Methods). Because cytoskeletal remodeling is closely linked to cell migration, we focused on cytoskeleton-related genes that were highly expressed in BC-seq-mediated scRNA-seq data and conducted transcription factor prediction. This analysis identified serum response factor (SRF) as a candidate TF (Methods, Fig. 3f).

SRF is a well-established regulator of actin cytoskeletal organization and focal adhesion assembly^34,35^. Based on these prediction results, we hypothesized that SRF is a TF activated by nuclear VEGFR2. Since transcription factor activity correlates with its binding to response elements, we used a Serum Response Element (SRE) reporter to measure SRF binding activity (Methods). Consistent with our hypothesis, SRF binding activity (bioluminescence signal) was increased under the SREBP2-OE condition, whereas it was reduced under the combined SREBP2-OE and VEGFR2-KD condition (Supplementary information, Fig. S8, left panel). To exclude the possibility that these changes were due to nuclear VEGFR2-induced alterations in SRF expression, we quantified SRF expression under both conditions and observed no significant differences (Supplementary information, Fig. S8, right panel). Taken together, these results reveal that nuclear VEGFR2 enhances cell migration by functioning as a transcriptional co-activator for SRF, thereby increasing SRF binding activity.

### Cholesterol trafficking via STARD4 to the cytoplasmic membrane is crucial for fast cellular migration

Having established that caveolae membrane microdomains are essential for caveolae-mediated endocytosis for VEGFR2 nuclear translocation, specifically mediated by the Cavin1 and Cavin3 trafficking proteins, we next investigated how increased cholesterol is transferred to the cytoplasmic membrane, a process critical for the initiation of caveolae formation. Upon examining cholesterol trafficking-related genes within our DEGs, we found that STARD4, a well-known mediator of intracellular cholesterol transport, was upregulated in FMCs (Fig. 4a).

**Figure 4.**
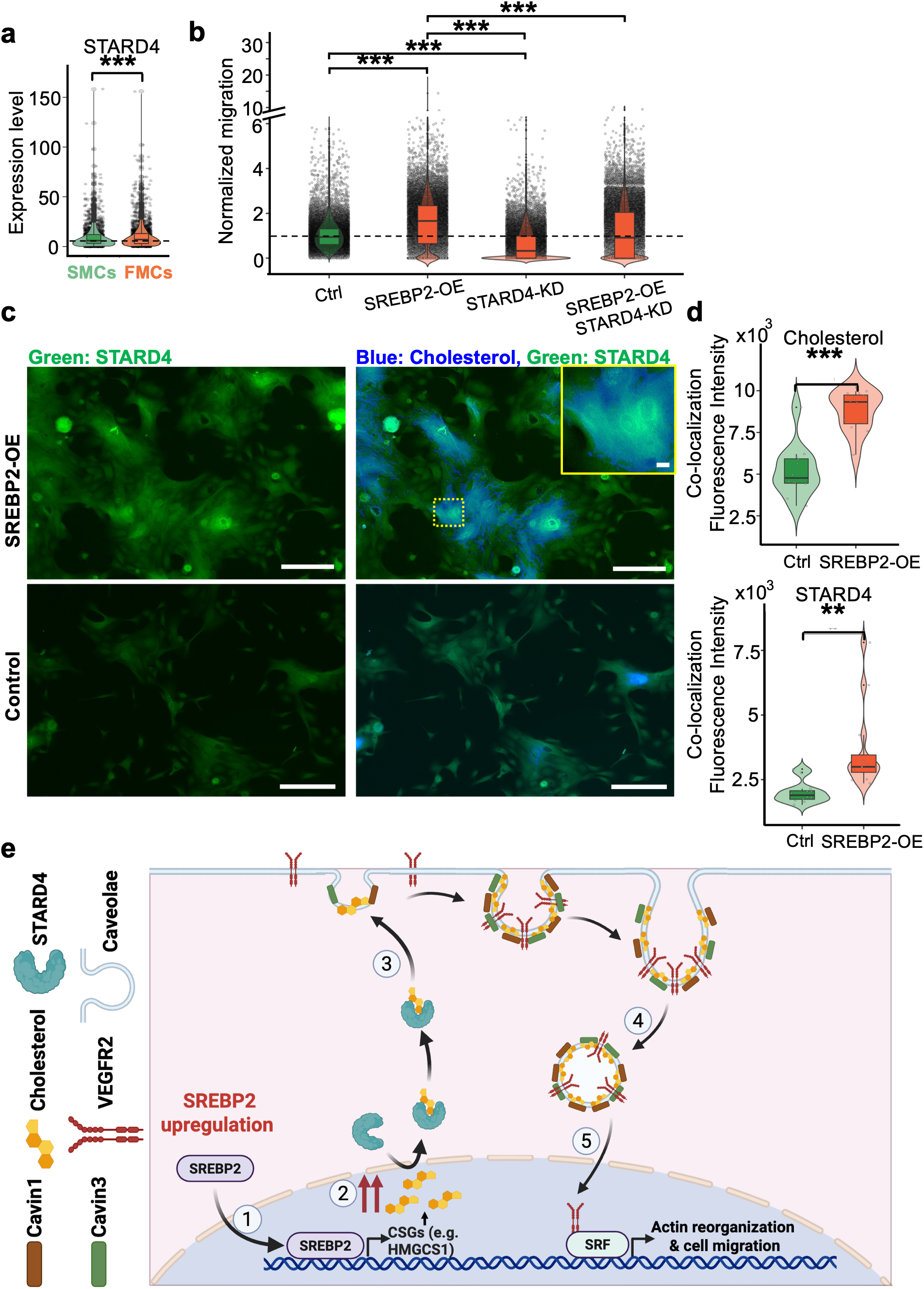
Cholesterol trafficking via STARD4 to the cytoplasmic membrane is crucial for fast cellular migration. **a.** A violin plot shows gene expression level of STARD4 for SMCs and FMCs. **b.** A violin plot shows normalized migration of cells under control, SREBP2-overexpression (OE), STARD4-knockdown (KD), or combined SREBP2-OE with STARD4-KD conditions. **c.** Representative immunofluorescence images of OSCC cells stained with STARD4 (green) and Cholesterol (blue) under control and SREBP2-OE conditions. Scale bars = 50 μm; 5 μm for insets. **d**. Fluorescence intensity of colocalized cholesterol (top panel) or STARD4 (bottom panel) signals under control (Ctrl) and SREBP2-OE conditions. **e**. Proposed mechanism: Elevated cholesterol, driven by SREBP2 (Step 1), is trafficked to cytoplasmic membranes by STARD4 (Step 2), initiating the formation of caveolae membrane microdomains (Step 3). This, in turn, enables Cavin 1/3-mediated endocytosis (Step 4) and VEGFR2 nuclear translocation, and activates VEGFR2-SRF-mediated actin reorganization (Step 5), ultimately leading to enhanced cell migration. For a, b, d, *P* values were calculated using the Wilcoxon rank sum test with Bonferroni correction.

We hypothesized that STARD4 plays a key role in transporting cholesterol to the membrane under SREBP2-upregulated conditions. This relationship was supported by a reduction in cellular migration when SREBP2 was overexpressed under the STARD4-knockdown (KD) condition, compared to the SREBP2-OE condition (Fig. 4b). Furthermore, a clear reduction in migration was observed in the STARD4-KD condition alone, indicating its essential role in facilitating cellular migration (Fig. 4b; The qPCR results for STARD4 gene expression after KD are shown in Supplementary information, Fig. S9). To further validate this mechanism, we performed co-immunostaining of STARD4 and cholesterol. We observed both increased STARD4 protein expression and elevated cholesterol levels under the SREBP2-OE condition (Fig. 4c, d, Supplementary information, Fig. S10). Most importantly, there was clear co-localization of higher STARD4 expression and cholesterol (Fig. 4d), confirming the role of STARD4 in trafficking cholesterol in cells.

Collectively, our findings reveal a mechanistic cascade (Fig. 4e) in which elevated cholesterol, driven by SREBP2, is trafficked to cytoplasmic membranes by STARD4, initiating the formation of caveolae membrane microdomains (Fig. 4e, Step 1-3). This, in turn, enables Cavin 1/3-mediated endocytosis (Fig. 4e, Step 4) and VEGFR2 nuclear translocation, and activates VEGFR2-SRF-mediated actin reorganization (Fig. 4e, Step 5), ultimately leading to enhanced cell migration or the transition of cells into a fast migratory state.

### A robust gene-expression signature derived from BC-seq predicts metastasis in OSCC patients

While studies have shown that cells exhibiting aggressive migratory behavior are closely linked to metastasis in both *in vitro* and *in vivo* models^36,37^, the molecular mechanisms underlying this behavior have not yet been directly correlated with metastatic status in clinical settings. To test our initial hypothesis—that the molecular mechanisms underpinning tumor metastasis are closely associated with RNA expression profiles that drive aggressive cellular migration—and to validate whether the temporal RNA expression identified in subpopulations of cells exhibiting transiently aggressive migratory behavior is clinically relevant, we identified a set of gene markers (Supplementary information, Fig. S11), that were consistently detected by two or more of the four analytical models analyzed. These models included supervised machine learning algorithms, logistic regression and linear support vector classification, as well as Limma and Wilcoxon rank-sum analyses (Methods). Using NanoString’s nCounter bulk-cell profiling assays, we quantified RNA transcript levels of the identified gene markers in a cohort of 81 OSCC clinicopathological formalin-fixed paraffin-embedded (FFPE) samples. This cohort included 38 cases without metastasis and 43 cases with distant metastasis, selected from a larger database of ∼4,000 HNSCC cases (collected between 2006 and 2022) based on defined inclusion criteria, such as primary tumors, treatment-naïve status, and the availability of high-quality FFPE samples.

We first performed univariate tests and identified 39 significant genes associated with metastatic samples (Methods; Supplementary Fig. S11, highlighted in green and red; Methods). These genes include those related to cholesterol synthesis and metabolism signaling^38^, such as STARD4, CAVIN1, INSIG1, DDIT3, and SEC24D — the first three of which were already identified and highlighted in our earlier analyses. We reason that Cavin1, but not Cavin3, was significantly identified in the bulk-cell profiling because Cavin1 is the most abundant protein in the cavin coat^32^. Additionally, SNX12, LGALS1, TPM1 and TPM4 were found to be involved in the partial epithelial-to-mesenchymal transition (p-EMT) program. Genes associated with VEGFA-VEGFR2 signaling^38^, such as HSPA1A (also known as HSP70-1A) and SPARC, were also identified and are known to contribute to cell migration and tumor metastasis^39,40^. Notably, many downstream effectors of cellular movement and migration, particularly those linked to actin cytoskeleton remodelling and focal adhesion signaling^38^, were also present in these 39 genes. These include COL1A1, COL1A2, ACTA2, FN1, ACTN1, TGFB1I1, FBLIM1, VASP, PPP1R18, CD99, RHOC, LASP1, RRAS, SNAI2, CDKN2A, MYH9, and PFN1.

These results align with our earlier findings on cellular and molecular mechanisms, in which actin cytoskeleton and focal adhesion pathways, along with cholesterol synthesis and metabolism, were shown to correlate with aggressive cell migration. The identification of these genes in our bulk-cell profiling of metastatic samples further validates the reliability of the BC-seq-profiling data. These results provide evidence linking the molecular mechanisms driving aggressive cellular migration to metastatic status in clinical settings. Moreover, they validate our hypothesis and underscore the efficacy of profiling selectively aggressive cellular subsets to identify metastasis-associated gene signatures.

Finally, using a logistic regression model with Elastic Net regularization (Methods), we identified a 15-gene signature for metastasis stratification and internally validated its performance through 50 iterations of bootstrap resampling (red-highlighted genes in Supplementary Fig. S11). Among the 10 upregulated genes in the signature are: VASP, CD99, RHOC, TGFB1I1, COL1A2, INSIG1, STARD4, SPARC, HSPA1A, and SNX12 (Fig. 5a). To assess the performance of the 15-gene signature set in stratifying patients into high-or low-risk groups for distant metastasis, we defined an optimal threshold by maximizing the Youden index^41^ and used the .632 rule for all bootstrap-estimated metrics^42^, resulting in a sensitivity of 82.0% and an accuracy of 83.6% (Fig. 5b, Methods). This significantly surpasses the predictive accuracy of current clinicopathological metrics (e.g., age and tumor size), which have been reported to range between 20% and 30% in previous studies^17,18^, as well as in our 81-patient validation cohort (Fig. 5c). Moreover, to externally validate our gene signature, we performed a similar analysis to calculate prediction scores using our 15-gene signature on patients with regional lymph node metastasis (Stage N2/N3) and without (Stage N0) from the TCGA database^43^, including both HNSCC patients and the OSCC patient subgroup (Supplementary information, Fig. S12). The prediction scores showed significant differences between N2/N3 and N0 patients in both TCGA-HNSCC and TCGA-OSCC cohorts, with the OSCC subgroup showing a particularly stronger statistical association (Supplementary information, Fig. S12). This result is consistent with the fact that the gene signature was originally derived from OSCC-specific profiling. These external validation results further confirm the predictive power of the signature and support its potential application as a clinical prognostic tool.

**Figure 5.**
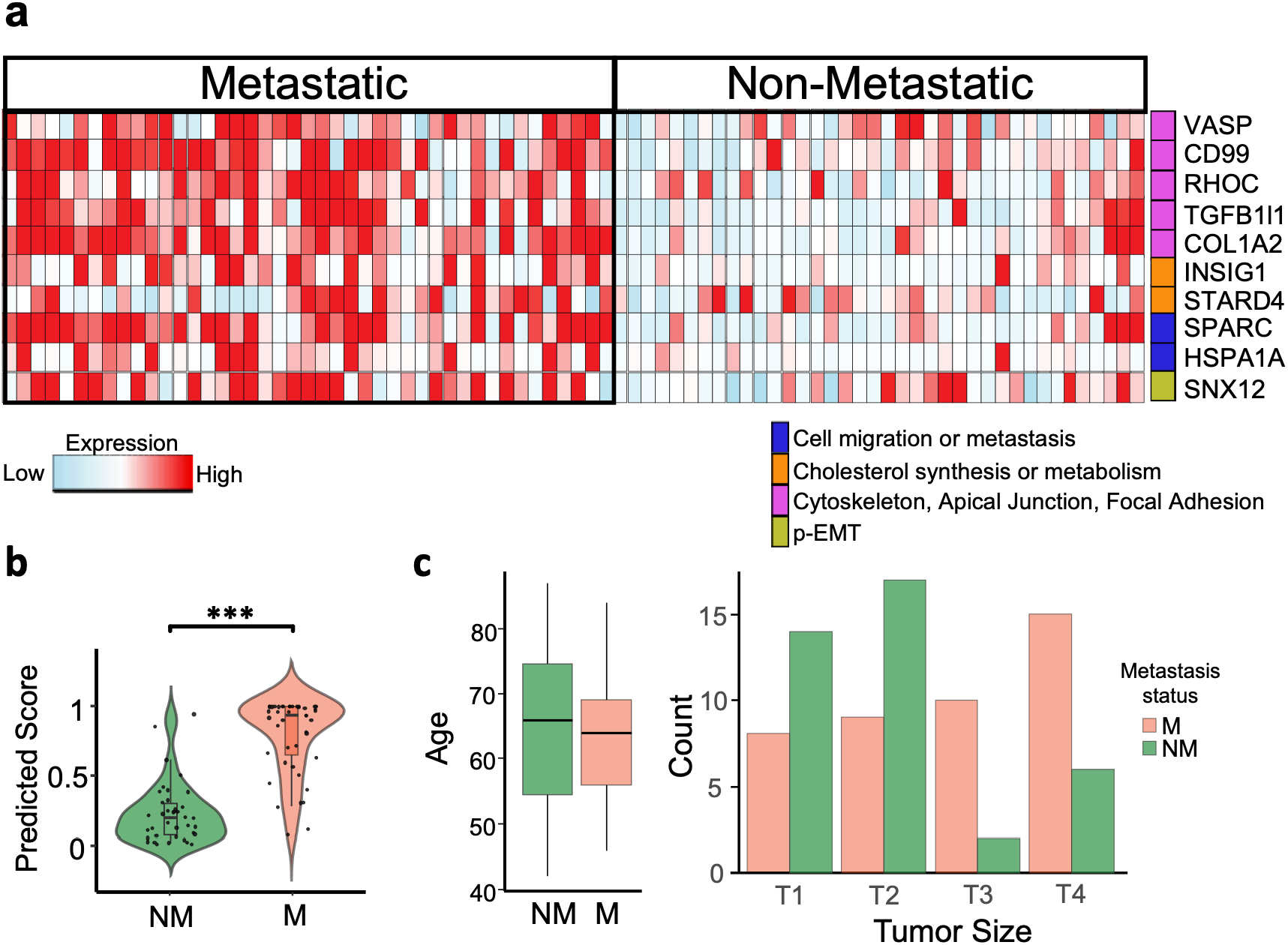
BC-seq-derived gene-expression signature predicts metastasis in OSCC patients. **a**. NanoString nCounter analysis of RNA transcript levels for the gene signatures (upregulated in the metastatic patients) in 81-OSCC clinicopathological samples (38 non-metastatic (NM) and 43 with distant metastasis (M)). Gene functions were color-coded as follows: blue indicates genes related to cell migration or tumor metastasis; orange indicates genes involved in cholesterol synthesis or metabolism; pink indicates genes related to the cytoskeleton and focal adhesion; and green indicates genes associated with partial EMT (p-EMT). **b**. Predicted scores of 43 M and 38 NM patient samples calculated using a logistic regression model with Elastic Net regularization. P values were calculated using the Wilcoxon rank sum test with Bonferroni correction. **c**. Clinicopathological data, specifically age (Left panel) and tumor size (Right panel), for the 81-patient validation cohort.

In conclusion, understanding and targeting metastatic dissemination remain challenging, in part because the molecular mechanisms governing aggressive tumor cell migration are difficult to resolve using conventional approaches. This challenge reflects the dynamic and transient nature of highly migratory cell states, which often arise within small subpopulations that are not readily captured by static molecular profiling. By directly linking migratory behavior to transcriptional state, our study demonstrates that resolving temporal gene expression associated with aggressive migration provides insight into both the biological and clinical relevance of these states. Using this behavior-guided framework, we identify a cholesterol-regulated signaling program involving the trafficking proteins STARD4 and CAVIN1/3 that contributes to aggressive migration in oral squamous cell carcinoma. In addition, we define migration-associated gene programs encompassing cholesterol metabolism, VEGFA-VEGFR2 signaling, and actin cytoskeletal regulation that are reflected in clinical outcomes. Together, these findings establish behavior-coupled transcriptomic profiling as an approach for dissecting the molecular basis of aggressive tumor cell migration and suggest a general strategy for uncovering migration-associated mechanisms and clinically informative biomarkers across tumor types.

## Materials and Methods

### HNSCC primary cultures for BC-seq assays

#### Ethics declarations

Patient-derived HNSCC tissues and data were used according to “The Code of Conduct for Responsible Use” and “The Code of Conduct for Health Research” as stated by the Federation of Dutch Medical Scientific Societies. The use of HNSCC samples was approved by Erasmus University Medical Center ethics committee (MEC-2022-0002). Informed consent was obtained from all patients for use of biopsy materials in research.

#### HNSCC patient-derived primary cultures for BC-seq

Primary tumors from 10 HPV-negative oral squamous cell carcinoma (OSCC) samples, the most common type of HNSCC, were used in the BC-seq assay. Please note that BC-seq enables the enrichment of selected cell types of interest — in this case, fast-migrating cells (FMCs) and slow-migrating cells (SMCs) — and allows direct linking of aggressive cell migration to transcriptomic profiles without interference from non-specific cells. As a result, cell type-or phenotype-specific profiles are greatly enhanced by BC-seq, and profiling a relatively small number of patient samples (> 5 samples, validated in the lab) is often sufficient to identify relevant molecular mechanisms.

After tumor excision, the tumor tissue was subjected to a commercial mechanical/enzymatic dissociation system (GentleMACS, Miltenyi Biotec, Bergisch Gladbach, Germany) according to the manufacturer’s protocol. Briefly, the tumor was cut into small fragments about 2-3 mm in length and put in a C-tube (Miltenyi Biotec) with clear DMEM/F12 medium (ThermoFisher Scientific) and solutions H, R and A (all from Miltenyi Biotec) according to the manufacturer’s recommendation; the digest mix containing the tumor was then subjected to three 36-sec mechanical disaggregation steps (program h_tumor_01, though tumors) in the GentleMACS dissociator interspersed by two 30-min rotating incubations (MACSmix Tube Rotator) at 37°C performed after the first and the second disaggregation steps, respectively. After disaggregation, dissociated cells were passed through 70 μm strainers (MACS SmartStrainers) and then cultured in a T25 culture flask (Green Bioresearch) coated with Cultrex basement membrane matrix (Bio-techne R&D systems, #3432-005-01) in the PNEU medium^44^ supplemented with PneumaCultTM-Ex Plus 50X Supplement (STEMCELL Technologies), 96 ng/mL of hydrocortisone solution, 1% of 200 mM L-Glutamine, 2.5 μg/mL of Amphotericin-B (ThermoFisher Scientific), 100 U/mL of Penicillin and 100 μg/mL of Streptomycin (Sigma-Aldrich). Cells were maintained in a humidified incubator (37°C, 5% CO2) and culture medium was refreshed every 3-4 days. Cultures were expanded after reaching ∼80-90% confluency to maintain a cell density of 200-800 cells/mm^2^. Propagation of cells was performed using enzymatic dissociation (1 mL Accutase, STEMCELL Technologies). Cells at early passages (∼5 passages) were used for the BC-seq assay.

### BC-seq pipeline and analysis

#### Platform overview

Behavior-coupled single-cell transcriptomic sequencing (BC-seq) platform operates as follows: i) The process begins by imaging a large number of single cells (∼10^4-5^), pre-incubated with photoactivatable dyes (see “Platform detail” below), using our Ultrawide Field-of-View Optical (UFO) Microscope ^45^. ii) The pipeline then processes these images in real-time, tracking individual cells to identify subpopulations in the fast (or slow)-migration state. iii) Next, it selectively and automatically photolabels cells of interest (a technique referred to as phototagging) using a 405 nm laser directed by photo-patterning devices (galvo mirrors) integrated into the microscope. iv) Finally, target cells are isolated through fluorescence-activated cell sorting (FACS) and proceed to single-cell (RNA) sequencing. This platform is largely similar to our original FUNseq pipeline^45^, with the modification of the behavior analysis (see below). We refer to this platform as behavior-coupled scRNA-seq to emphasize its ability to profile single cells based on cellular behaviors.

#### Platform detail

The day before conducting cell migration assays, OSCC cells were seeded at a density of approximately 300-500 cells/mm² in the micro-wells of glass-bottom dishes (Cellvis), pre-coated with Cultrex (Bio-techne R&D systems, #3432-005-01). After allowing the cells to adhere for 4 hours, ensuring complete attachment, the medium was increased to 2 mL, and a nuclear stain—SPY650-DNA (Spirochrome) (1:2000 dilution, λ_ex_: 652 nm, λ_em_: 674 nm)—was then added and incubated overnight. Prior to imaging, cells were incubated with 30 μM of photoactivatable dye (Janelia Fluor 549, TOCRIS) for 10-20 minutes. Imaging was conducted within a humidified incubator (37°C, 5% CO2) using our UFO Microscope^45^. Cell nuclei were imaged with a 637 nm laser at 4-minute intervals, followed by real-time nuclear segmentation and tracking of cell locations throughout the imaging period, as described in Chou et al.^46^. After one hour of imaging and real-time analysis (∼5 minutes post data acquisition), target cells (such as FMCs) were identified and phototagged using a 405 nm laser (100 J/cm²) through a photopatterning device (galvo mirrors implemented in the UFO system). The signal of photoactivation (λ_ex_: 549 nm, λ_em_: 571 nm) was visualized at 532 nm, where non-phototagged cells remained dark. Following phototagging, cells were dissociated using 200 μL Accutase (STEMCELL Technologies) enzymatic treatment, and resuspended in Hank’s Balanced Salt Solution (HBSS) to achieve a single-cell suspension. This suspension was then transferred to a BD FACSMelody sorter (BD Biosciences), where phototagged target cells were identified and sorted based on both the nuclear stain signal and the phototagging signal. Single cells were sorted into 384-well plates, which were subsequently centrifuged and stored at -80°C until further processing.

#### Identification and characterization of cellular behaviors, namely fast- and slow-migrating cells (FMCs and SMCs)

UFO-acquired images were processed for individual cell (nuclei) segmentation, tracking and (migratory) trajectory analysis:

1. Individual cells in each frame were segmented using ground-truth-assisted trainable Weka segmentation (GTWeka)^46^, a machine learning-based method capable of rapidly (within 5 seconds) and accurately identifying and separating approximately 20,000 individual cells per frame. Each cell is characterized by features such as location, morphology, and brightness. GTWeka is designed to be user-friendly, requiring minimal data annotation for training, making it accessible even to users without prior experience in training deep learning models. The annotation process is straightforward, guided by simple instructions^46^.
2. We subsequently utilized the modified Tracking with Gaussian Mixture Model (mTGMM)^46^ to monitor cell locations over time, linking them into continuous time sequences^46^. For each individual cell, the connection between two consecutive frames, *t* and *t+1*, was established based on the similarity of their features (location, morphology, and brightness) across these adjacent frames. For instance, the cell’s positions at two consecutive time points were required to exhibit a nearest-neighbor relationship. The calculations for all cells within a frame (∼20,000 cells) were performed efficiently, taking less than one minute, using the linear assignment method. This process enabled the generation of a migratory trajectory for each cell across all frames.
3. To identify cells in fast or slow migratory states, we calculated the speed of each cell from time-lapse movies and ranked them in descending order. Cells in the fast or slow migratory states were defined as those whose migratory speeds fell within the top or bottom 5% of the speed histogram distribution. Additionally, we used Mean Squared Displacement (MSD) analysis (Supplementary Fig. S13) to characterize cell migration patterns, distinguishing between directional and non-directional movement. The MSD was calculated using the following equation: 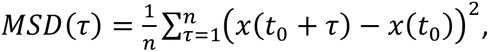 where *x*(*t_0_*) represents a cell’s location at time point *t₀*, typically the initial time point. The MSD considers various time intervals as τ progresses from 1 to *n*, where *n* is approximately half the total number of time points (*T/2*). The MSD plot for FMCs revealed that these cells exhibit a clear pattern of directional movement.

#### BC-seq-mediated single-cell RNA sequencing and analysis

FMCs and SMCs processed through the BC-seq pipeline were sorted via FACS into 384-well plates pre-filled with barcoded primers.

These plates were subsequently frozen followed by single-cell RNA library preparation and sequencing (150,000 reads/cell) using the SORT-seq procedures^47^. In brief, mRNA from each cell was reverse-transcribed into complementary double-stranded DNA (cDNA). Subsequently, all contents from each well were pooled, followed by in vitro transcription, library preparation, and Illumina sequencing, carried out according to the protocol outlined by Muraro et al.^47^.

Raw FASTQ files from 10 OSCC patient sequencing were aligned to the reference genome (Homo sapiens, Hg38) using the Spliced Transcripts Alignment to a Reference (STAR) aligner (v2.7.10)^48^, with the STARsolo workflow to generate gene count matrices. These counts were then corrected using a Poisson model^49^. These data underwent processing using the Seurat package (v5.1.0)) ^50^ in R. Quality control (QC) measures were implemented to exclude low-quality cells based on specific criteria. Cells with mitochondrial ratios exceeding 35%, unique molecular identifier (UMI) counts below 2,000, feature counts under 2,000, or potential doublets identified by gene counts above 15,000 were removed from the analysis. After QC, 1405 OSCC cells were used for downstream analyses in this study.

Standard Seurat (v.4.4.0) procedures were employed, including log-normalization^51^, scaling, and the selection of the top 10,000 most variable genes for dimensionality reduction analyses (e.g., PCA and UMAP plots). Seurat-based cell clusters were generated using the original Louvain algorithm at the default resolution of 0.7. Module scores were calculated using the ‘AddModuleScore’ function in the Seurat package^52^, incorporating specific gene signatures (e.g., cholesterol synthesis and metabolism^53^ and actin cytoskeleton signaling-related genes^38^). To assess the significance of module scores and gene expression levels within each group, Wilcoxon signed-rank tests were performed, with Bonferroni correction applied for multiple comparisons.

Cell types were identified using the R package SingleR^54^. The BlueprintEncodeData dataset t^55^, a built-in reference in SingleR, was employed for all annotations. SingleR generates cell type annotations at the single-cell level by producing an assignment score matrix, where each cell is assigned a score for each cell type label from the reference. To convert these single-cell annotations to cell-cluster level annotations, we assigned the reference label with the highest assignment score to each cell, based on the default parameters. Fewer than 1% of non-epithelial cells were detected and excluded prior to downstream analysis.

Differential gene expression analysis was performed using the Linear Models for Microarray Data (limma) package (v.3.54.2))^56^, with statistical significance determined by adjusting p-values using the Benjamini-Hochberg correction. Genes with an adjusted p-value less than 0.1 and a log2 fold change greater than or equal to 0.1 were considered significantly upregulated. Pathway Enrichment Analysis was subsequently conducted on these significant genes using the Enrichr tool^57^, leveraging the WikiPathways 2024 database^38^.

### Identification of gene markers associated with metastasis using multiple models

Gene markers for metastasis prediction in OSCC patients were identified based on their consistent detection by two or more of the four analytical models analysed: Wilcoxon rank sum (Seurat package (v.4.4.0)), Limma (v.3.54.2) and two supervised machine learning models, logistic regression (LR) and linear support vector classification (LSVC) with the Python package Scikit-learn (version 1.0.2). Differentially expressed genes (DEGs) identified by Limma are described in the “Single-cell RNA sequencing and analysis” section, while those identified by LR and LSVC were derived as follows:

Analyses were conducted in Python (version 7.2.2). Doublets were removed, and good QC cells were selected based on criteria similar to those in the “Single-cell RNA sequencing and analysis” section. Cells with feature counts between 4,000 and 10,000 were included for downstream analysis. Filtered datasets were log-normalized, and batch effects across samples were corrected using Combat (version 0.3.3). After normalization and batch effect correction, gene signatures distinguishing FMCs from SMCs were selected based on weights exceeding ±0.15 in either the LR or LSVC model. Finally, a Venn diagram was used to identify genes that overlapped across four models (Wilcoxon, Limma, LR, and LSVC). Those overlapping (two or more models) and highly ranked genes were designated as our top pre-candidate gene signatures for the nCounter assay (Supplementary Fig. S6).

### NanoString nCounter gene expression assays

#### RNA isolation and quality control

The assay was performed on formalin-fixed paraffin-embedded (FFPE) samples from the Erasmus University Medical Center HNSCC cohort. Specifically, an 81-patient cohort with OSCC patient samples, including cases with no metastasis and with distant metastasis, selected from a larger database of ∼4,000 HNSCC cases (covering the years 2006-2020) based on defined inclusion criteria (e.g., primary tumors, treatment-naïve status, availability of high-quality FFPE samples). The cohort comprises 43 distant-metastatic samples and 38 non-metastatic samples. Tumor-rich areas were first identified on an H&E slide by HNSCC pathologists and then macrodissected into 20 to 30 unstained slices (∼10 μm each) using a xylene-based deparaffinization method. Total RNA was extracted using the Qiagen RNeasy FFPE kit (Qiagen), following the manufacturer’s instructions. The RNA concentration and quality of all samples were assessed using the Bioanalyzer PICO RNA kit (Agilent).

#### nCounter assay

The nCounter® Max Analysis System (NanoString Technologies) was used to perform the assay following the manufacturer’s standard protocol. Custom CodeSets, consisting of pre-built capture probes tagged with biotin and reporter probes labeled with fluorescently color-coded molecular barcodes, were designed and constructed by NanoString Technologies for selected gene sets (gene signatures related to metastasis prediction as previously described).

Approximately 300 ng of total mRNA (60 ng/µL) extracted from FFPE tissues was used for the assay. Hybridization reactions were prepared according to the manufacturer’s instructions and incubated for 18 hours at 65°C using Standard CodeSets reagents. Following hybridization, samples were processed using the nCounter Prep Station and immobilized onto the sample cartridge for data quantification, which was performed using the nCounter Digital Analyzer (NanoString Technologies). The nCounter Prep Station and Digital Analyzer together form the nCounter® Max Analysis System.

#### Processing and quality control of nCounter data

Data quality from the NanoString assay was assessed, and normalization was performed using the nSolver Analysis Software v3.0 (NanoString Technologies). Initially, raw counts were corrected for background noise by calculating geometric means of 8 negative control probes. Subsequently, counts were further corrected using geometric means of 6 internal positive control spike-ins in each lane/sample to correct potential sources of variation across the samples. These negative and positive control probes are built into the nCounter assay protocol. To adjust for variations in RNA input volume, expression levels were normalized using the geNorm-selected housekeeping genes (ALAS1, POLR2A, TBP)), a feature integrated into the nSolver software. The final normalized count data were then used for subsequent significance analysis.

### Univariate tests to identify 39 significant genes

To identify genes associated with metastasis, we performed univariate tests comparing M (Metastasis) vs. NM (No-Metastasis) for each gene. Depending on distributional assumptions, we used the Shapiro-Wilk normality test to determine whether the two groups (Metastasis and No-Metastasis) followed a normal distribution. To assess the equality of variances, we performed the Fisher-Snedecor test. If the data were normally distributed with equal variances, an unpaired two-sample t-test was conducted. The t-test was unpaired as the samples originated from different patients, making the groups independent. Conversely, if the data were normally distributed but exhibited unequal variances, the Welch’s t-test was applied. For data that were not normally distributed, we utilized the non-parametric Wilcoxon rank-sum test (Mann-Whitney U test), which does not assume equal variances. 39 genes passed the pre-specified significance threshold and we take them in consideration before the multivariate modelling using a logistic regression model.

### 15-gene signature identification and metastasis risk prediction

To identify a robust gene signature set for predicting metastatic status (Metastasis vs. No-Metastasis), we developed and internally validated a logistic regression model. The input dataset consisted of an independent cohort of 81 OSCC patient samples, including cases with no metastasis and with distant metastasis. In this cohort 100 genes were measured using the nCounter Nanostring platform. Internal validation was performed through 50 iterations of bootstrap resampling with replacement. In the bootstrap analysis, we used 63% of unique samples for training and the remaining 37% for testing. In each iteration, we developed a logistic regression model with regularization (Elastic Net) using the *glmnet* package (v4.1-8) in R (v4.4.0). In the Elastic Net model, the regularization parameter lambda was selected via cross-validation by choosing *lambda.1se*. The mixing parameter alpha was selected based on the Area Under the Curve (AUC) of the Receiver Operating Characteristic (ROC) curve. In each bootstrap iteration, we evaluated the AUC and summarized the performance using the .632 bootstrap correction method to account for resampling bias^42^. For threshold-based metrics derived from the confusion matrix (sensitivity and accuracy), the optimal threshold was determined using the Youden index^41^, implemented with the *pROC* R package (v1.18.5). The final model, consisting of 15 genes (Supplementary Fig. S6), was developed by applying the same pipeline used in the bootstrap iterations to the entire dataset.

To externally validate our gene signature, we performed a similar analysis to calculate prediction scores using our 15-gene signature on patients with regional lymph node metastasis (Stage N2/N3) and without (Stage N0) from the TCGA database^43^, including both HNSCC patients and the OSCC patient subgroup. In the TCGA-HNSCC and -OSCC datasets, log₂-transformed FPKM-UQ values (log₂[FPKM-UQ + 1]) were extracted for these 15 genes, and a sample-wise risk score was computed as the weighted sum of expression values using the coefficients obtained from the trained model. Scores were min–max normalized to the [0, 1] range for interpretability and visualization. The score distributions were assessed using the Wilcoxon rank-sum test.

### Identification of serum response factor (SRF) as the transcription factor driving cell migration

To identify potential transcription factors regulating actin cytoskeleton-related genes, we first performed KEGG pathway enrichment analysis on the differentially expressed genes obtained from BC-seq-mediated scRNA-seq data. Genes annotated to the “regulation of actin cytoskeleton” pathways were selected as the input gene set. Transcription factor (TF) prediction was then carried out using the **RcisTarget** R package (version 1.24.0)^33^, a tool designed for TF motif enrichment analysis. The gene-motif ranking database *hg19-500bp-upstream-7species.mc9nr.genes_vs_motifs.ranking.feather* was used for motif enrichment analysis. RcisTarget tests whether TF motifs are significantly enriched in the regulatory regions of the input genes and provides candidate TFs with evidence of motif enrichment, from which the likely upstream regulators can be identified. SRF was identified as the sole TF after the analysis.

### Validation assays

#### Cholesterol depletion and enrichment assays

Cholesterol depletion and enrichment assays were conducted in triplicate using methyl-β-cyclodextrin (MβCD) (Sigma-Aldrich) for depletion and cholesterol-MβCD complex for enrichment. MβCD was dissolved in distilled water. To prepare the cholesterol-MβCD complexes, cholesterol (Sigma-Aldrich) dissolved in a methanol-chloroform mixture (2:1, v/v) (both from Thermo Fisher Scientific) was added to a 5% (w/v) MβCD solution, which was maintained at 80 °C in a water bath. The mixture was stirred at 80 °C until the initially precipitated cholesterol was fully dissolved.

For the cholesterol depletion assay, cells were initially treated with 10 mM MβCD for one hour, then washed three times with DPBS before fresh medium was added to the culture ^58^. MβCD depletes cholesterol by encapsulating hydrophobic components of the plasma membrane within its inner hydrophobic cavity, continuously extracting cholesterol from the outer leaflet of the membrane^59^. Cholesterol enrichment was carried out by first treating the cells with 10 mM MβCD for one hour, followed by incubation in fresh medium containing 100 μM cholesterol-MβCD complexes for two hours. After treatment, the cells were imaged, and migration analysis was performed as previously described. To account for potential variations between assayed cell batches, the migration speeds for both the control and experimental conditions, assessed concurrently, were normalized to the average migration speed of the control group.

### SREBP2 signaling, CAVIN1, CAVIN3, CAV1, VEGFR2 and STARD4 inhibition assays

To suppress the expression of the indicated target genes, we designed short hairpin RNA (shRNA) constructs specific to each transcript. Each shRNA contained a 21-nucleotide stem separated by a loop sequence, enabling the formation of a hairpin structure that is efficiently processed by the endogenous RNA interference pathway.

### Preparation of anti-SREBP2 shRNA insert oligonucleotides

To inhibit SREBP2 signaling, we designed short hairpin RNA (shRNA) constructs targeting the SREBP2 transcript. shRNA targeting sequence was selected as follows:

Forward: 5’-CCGGGACCTGAAGATCGAGGACTTTCTCGAGAAAGTCCTCGATCTTCAGGTCTTTTTG-3’ Reverse: 5’-AATTCAAAAA GACCTGAAGATCGAGGACTTTCTCGAG AAAGTCCTCGATCTTCAGGTC-3’

### Preparation of anti-CAVIN1 shRNA insert oligonucleotides

To inhibit CAVIN1 gene, we designed shRNA constructs targeting the CAVIN1 transcript. shRNA targeting sequence was used as follows:

Forward: 5’-CCGGGTGGAGGTTGAGGAGGTTATTCTCGAGAATAACCTCCTCAACCTCCACTTTTTG-3’ Reverse: 5’-AATTCAAAAAGTGGAGGTTGAGGAGGTTATTCTCGAGAATAACCTCCTCAACCTCCAC-3’

### Preparation of anti-CAVIN3 shRNA insert oligonucleotides

To inhibit CAVIN3 gene, we designed shRNA constructs targeting the CAVIN3 transcript. shRNA targeting sequence was used as follows:

Forward: 5’-CCGGGCCAGCGCTTTCCAGAAGGCACTCGAGTGCCTTCTGGAAAGCGCTGGCTTTTTG-3’ Reverse: 5’-AATTCAAAAAGCCAGCGCTTTCCAGAAGGCACTCGAGTGCCTTCTGGAAAGCGCTGGC-3’

### Preparation of anti-CAV1 shRNA insert oligonucleotides

To inhibit CAV1 gene, we designed shRNA constructs targeting the CAV1 transcript. shRNA targeting sequence was used as follows:

Forward: 5’-CCGGGACGTGGTCAAGATTGACTTTCTCGAGAAAGTCAATCTTGACCACGTCTTTTTG-3’ Reverse: 5’-AATTCAAAAAGACGTGGTCAAGATTGACTTTCTCGAGAAAGTCAATCTTGACCACGTC-3’

### Preparation of anti-VEGFR2 shRNA insert oligonucleotides

To inhibit VEGFR2 gene, we designed shRNA constructs targeting the VEGFR2 transcript. shRNA targeting sequence was used as follows:

Forward: 5’-CCGGGCGGCACGAAATATCCTCTTACTCGAGTAAGAGGATATTTCGTGCCGCTTTTTG-3’ Reverse: 5’-AATTCAAAAAGCGGCACGAAATATCCTCTTACTCGAGTAAGAGGATATTTCGTGCCGC-3’

### Preparation of anti-STARD4 shRNA insert oligonucleotides

To inhibit STARD4 gene, we designed shRNA constructs targeting the STARD4 transcript. shRNA targeting sequence was used as follows:

Forward: 5’-CCGGTGACCTTGTCTATAGTATAATCTCGAGATTATACTATAGACAAGGTCATTTTTG-3’ Reverse: 5’-AATTCAAAAATGACCTTGTCTATAGTATAATCTCGAGATTATACTATAGACAAGGTCA-3’

These sequences were used to design complementary oligonucleotides compatible with the third-generation pLKO.1-TRC lentiviral transfer vector (Addgene #10878). The oligonucleotides were synthesized (Integrated DNA Technologies) with appropriate overhangs for directional cloning into the AgeI and EcoRI sites of pLKO.1-TRC. Oligonucleotides were annealed by mixing equal molar amounts (100 μM, 6 μL each) in 3 μL 10x NEB buffer 2 (10mM TrisHCl, 50mM NaCl, 10mM MgCl2, 1mM DTT, all from Thermo Fisher Scientific) and 15 μL nuclease-free water (Qiagen) (total volume 30 μL), heating to 95°C for 5 minutes, and slowly cooling to room temperature. The annealed oligonucleotides were then ligated into AgeI/EcoRI-digested pLKO.1-TRC vector using T4 DNA ligase (New England Biolabs) according to the manufacturer’s protocol. Ligated plasmids were transformed into DH5α competent *E. coli* (Thermo Fisher Scientific) and positive clones were identified by colony PCR and confirmed by Sanger sequencing. As a negative control to assess non-specific effects, we used an empty pLKO.1-puro lentiviral plasmid containing a non-coding insert (Addgene #8453). Both interfering shRNA and control plasmids contained a puromycin resistance gene as a selectable marker for downstream selection of stably transduced cells. Plasmid DNA was purified using an endotoxin-free plasmid Miniprep kit (Qiagen) according to the manufacturer’s instructions. DNA concentration and purity were assessed using a DeNovix spectrophotometer (DeNovix). This approach allows for the specific knockdown of the gene expression to study its signaling pathway, with appropriate controls to account for non-specific effects of lentiviral transduction and shRNA expression.

#### SREBP2 and VEGFR2 overexpression assays

To overexpress SREBP2 and VEGFR2, we used a lentiviral vector system. The transcription factor open reading frame (ORF) for SREBP2 was obtained from the TFORF2861 plasmid (Addgene #143644) and VEGFR2 expression vector was obtained from pHAGE-KDR plasmid (Addgene #116754). These plasmids contained a puromycin resistance gene as a selectable marker for stable transduction. Lentiviral particles were produced as described in the lentivirus production section. Imaging and migration assessment were performed in triplicate. Confirmation of SREBP2 and VEGFR2 gene overexpression was performed through qPCR.

Following treatment with SREBP2-OE, VEGFR2-OE and anti-SREBP2, -CAVIN1, -CAVIN3, -CAV1, -VEGFR2 and STARD4 shRNA viruses as well as the control viruses (plasmid backbone only), cells were imaged, and migration analyses were conducted in accordance with previously described. To account for potential variations between assayed cell batches, the migration speeds for both the control and experimental conditions, assessed concurrently, were normalized to the average migration speed of the control group.

#### Lentivirus production

Lentiviral particles were produced using HEK293 cells. One day prior to transfection, HEK293 cells were seeded in 10 cm dishes at approximately 50% confluency. Transfection was performed when cells reached 70-80% confluency, typically 24 hours after seeding. For each 10 cm dish, a plasmid mixture was prepared containing 8.5 μg of the vector of interest, 4.5 μg of psPAX2 (a second-generation lentiviral packaging plasmid, Addgene #12260), and 4.5 μg of pMD2.G (a VSV-G envelope-expressing plasmid, Addgene #14888). Transfection was carried out using Lipofectamine 3000 reagent (Thermo Fisher Scientific) according to the manufacturer’s instructions. Briefly, 53 μL of Lipofectamine 3000 reagent and 35 μL of P3000 Enhancer were combined with the plasmid mixture in 2.5 mL of Opti-MEM medium (Gibco). Viral supernatant was collected at 24 and 48 hours post-transfection. The collected medium was filtered through a 0.45 μm PVDF filter (Sigma-Aldrich) to remove cellular debris. Viral particles were then concentrated by ultracentrifugation using a Beckman Coulter ultracentrifuge. The concentrated virus was aliquoted and stored at −80 °C until use.

#### Lentivirus titration

Lentiviral titers were determined by infection of HeLa cells followed by antibiotic selection and colony counting. On day 1, HeLa cells were seeded in 6-well plates at 2 × 10^5^ cells per well. Serial dilutions of virus were added to the medium containing 10 μg/mL polybrene (Millipore) and incubated overnight at 37°C and 5% CO2. The culture medium was refreshed 24 hours post-infection. At 48 hours post-infection, the medium was replaced with 2 mL of culture medium containing 1 μg/mL puromycin (concentration determined from HeLa cell kill curve). Selective medium was renewed every 2-3 days. After completion of antibiotic selection, as indicated by complete cell death in the control well, cells were washed once with DPBS (Gibco) and stained with 1% crystal violet solution (1% w/v in 10% ethanol, Sigma-Aldrich) for 15 minutes. Wells were then washed several times with DPBS. Stained colonies were counted using an automated gel counter (GelCount, Oxford Optronix). Viral titer in transducing units per mL (TU/mL) was calculated using the following formula: TU/mL = (Number of colonies x Dilution factor) / Volume of virus (mL).

#### Lentivirus transduction

OSCC cells were seeded in 6-well plates at a density of 60% confluency. After 24 hours, the culture medium was replaced with fresh medium supplemented with 10 μg/mL Polybrene (Millipore). Lentiviral particles were added to the cells at a multiplicity of infection of approximately 6.25 × 10^4^ transducing units (TU) per mL. Following a 24-hour incubation period, the virus-containing medium was replaced with fresh culture medium. Puromycin selection was initiated 48 hours post-transduction using 1.5 μg/mL Puromycin (Thermo Fisher Scientific), a concentration predetermined by kill curve analysis for OSCC cells. Selection was maintained until complete cell death was observed in non-transduced control wells. Successfully transduced and selected OSCC cells were harvested using 1 mL Accutase (STEMCELL Technologies) and re-seeded onto Cultrex-coated (Bio-techne R&D systems, #3432-005-01) glass-bottom dishes 24 hours prior to imaging and migration assessment as previously described. All migration inhibition and activation assays following knockdown and overexpression were conducted in triplicate. Confirmation of target gene knockdown and overexpression were done via qPCR.

### Immunofluorescence assays

#### Cell Fixation and Preparation

OSCC cells (both SREBP2-overexpressing and control cells) were fixed in 4% paraformaldehyde (PFA; methanol-free, Fisher Scientific) diluted in DPBS (Gibco) for 15 minutes at room temperature. Cells were then permeabilized by two 10-minute treatments with 0.1% Triton X-100 (Sigma-Aldrich) in DPBS. The samples were washed with a blocking buffer consisting of 0.15% (w/v) glycine (analytical grade, Sigma-Aldrich) and 0.5% (w/v) bovine serum albumin (BSA; Sigma-Aldrich) in DPBS to minimize nonspecific binding.

#### Cavin 3 and VEGFR2 Double Immunostaining

The first primary antibody against Cavin 3 (PRKCDBP Polyclonal Rabbit Antibody, Proteintech, #16250-1-AP, 1:100) was added and incubated for 90 minutes. A species-specific secondary antibody (Goat anti-Rabbit IgG, Alexa Fluor™ 488, Invitrogen, #A-11008, 1:1000) was then applied and incubated for 1 hour. Subsequently, the second primary antibody targeting VEGFR2 (VEGFR2/KDR/Flk-1 Mouse Antibody, R&D Systems, #MAB3571-SP, 1:50) was added and incubated for 90 minutes, followed by a species-specific secondary antibody (Goat anti-Mouse IgG, Alexa Fluor™ 594, Invitrogen, #A-11037, 1:1000) and another 1-hour incubation.

#### STARD4 immunostaining

The first primary antibody against STRAD4 (STARD4 Polyclonal Rabbit Antibody, Thermo Fisher Scientific, #PA5-61172, 1:100) was added and incubated for 90 minutes. A species-specific secondary antibody (Goat anti-Rabbit IgG, Alexa Fluor™ 488, Invitrogen, #A-11008, 1:1000) was then applied and incubated for 1 hour.

All antibody incubations were performed under humid and dark conditions at room temperature. The permeabilization and blocking steps, as previously described, were repeated between each antibody incubation to ensure optimal staining. After immunostaining, cells were washed twice with 0.1% Triton X-100 and once with DPBS. A 20 μL mixture of DAPI, DAPCO, and Vectashield (1:1:1; Vectashield mounting medium, Vector Laboratories) was applied to the cells. A coverslip was placed and sealed with nail polish. Samples were immediately analyzed using a fluorescence microscope (near-infrared wide-field fluorescence Zeiss).

### Filipin staining for cholesterol detection

Filipin is a naturally fluorescent polyene antibiotic (extracted from *Streptomyces filipinensis*) that specifically binds to free cholesterol in cell, allowing visualization of cholesterol distribution by fluorescence microscopy using UV excitation.

To perform the staining, OSCC cells were cultured for overnight. After treatment (i.e. SREBP2 overexpression as described before) cells were rinsed 2-3 times with DPBS to remove culture medium. Cells were then fixed with 3% paraformaldehyde (Fisher Scientific) in DPBS at room temperature for 15 minutes. Following fixation, cells were rinsed again 2-3 times with PBS. To quench residual aldehydes from fixation, cells were incubated with 1.5 mg/mL glycine (analytical grade, Sigma-Aldrich) in DPBS for 10 minutes at room temperature. Cells were then stained with filipin working solution (Filipin complex ready-made solution, Sigma-Aldrich) (0.05 mg/mL filipin complex in PBS containing 10% fetal bovine serum) prepared fresh and protected from light, for around 1 hour at room temperature in the dark.

After staining, cells were washed 2-3 times with DPBS to remove unbound filipin. Stained cells were immediately analyzed by fluorescence microscopy using a UV filter set with excitation at ∼340-380 nm and emission at 385-470 nm to detect filipin-cholesterol complexes in SREBP2-overexpressing and control samples.

### SRF bioluminescence assay

SRE (Serum Response Element) Luciferase Reporter Lentiviruses (bpsbioscience) were used to transduce patient-derived OSCC cells, which were subsequently selected with puromycin to establish a stable SRE-luciferase reporter cell line. These cells were seeded into three wells of a 6-well plate and re-infected according to three conditions: Control, SREBP2-overexpression, and SREBP2-overexpression with VEGFR2-knockdown. After a second round of antibiotic selection, 10,000 cells from each condition were seeded per well (6 replicates per condition) in an opaque 96-well plate (Thermo Fisher Scientific), incubated overnight at 37°C and 5% CO_2_. The following day, cells were treated with 100 μL of luciferase substrate (Luciferase Reporter Assay Substrate Kit – Firefly, ab228530), incubated at room temperature in the dark for 20 minutes, and then the luciferase activity (i.e. SRF activity) was immediately monitored by measuring the bioluminescence signal intensity (relative light unites, RLU) using a SpectraMax iD3 luminometer.

### Confirmation of target gene (SREBP2, CAVIN1, CAVIN3, CAV1, VEGFR2, and STARD4) knockdown and overexpression via quantitative real-time PCR (qPCR)

#### RNA extraction

The assayed cells were detached using 200 μL Accutase (STEMCELL Technologies), and their total RNA was extracted using the RNeasy Micro Kit (Qiagen), following the manufacturer’s protocol. Briefly, cells were lysed in 350 μL RLT buffer (Qiagen) supplemented with 3.5 μL β-mercaptoethanol (Thermo Fisher Scientific). In the presence of an equal volume of 70% v/v Ethanol (VWR), the total RNA was then bound to the RNA extraction column (Qiagen).

The column was washed with 350 μL RW1 buffer (Qiagen) to remove unwanted biomolecules, and residual DNA was removed with an on-column DNase digestion. After DNase treatment (80 μL of DNase solution; 10 μL DNase plus 70 μL RDD buffer, Qiagen), any remaining salts were washed away with 500 μL RPE buffer (Qiagen). Finally, the purified total RNA was eluted in 16 μL RNase-free double-distilled water (Qiagen).

#### cDNA synthesis

cDNA was synthesized using the iScript cDNA Synthesis Kit (Bio-Rad). The reaction mixture was prepared by combining 500 ng of template RNA, 4 μL 5x iScript Reaction Mix (containing random hexamers and dNTPs), and 1 μL iScript Reverse Transcriptase in nuclease-free water (total volume of 20 μL). The reaction conditions were as follows: priming at 25 °C for 5 minutes, reverse transcription at 46 °C for 20 minutes, and enzyme inactivation at 95 °C for 1 minute in a Bio-Rad thermal cycler. To control for potential genomic DNA contamination in subsequent qPCR experiments, a no-RT control reaction, excluding the reverse transcriptase, was included as a negative control.

#### qPCR assay

Total RNA was extracted and reverse-transcribed to cDNA as previously described. SREBP2, CAVIN1, CAVIN3, CAV1, VEGFR2 and STARD4 mRNA expressions were quantified using qRT-PCR with SsoAdvanced Universal SYBR Green Supermix on a CFX96 Real-Time PCR Detection System (Bio-Rad). Gene-specific primers were designed to span exon-exon junctions:

SREBP2 primers: 5ʹ-AACGGTCATTCACCCAGGTC-3ʹ (forward), 5ʹ-GGCTGAAGAATAGGAGTTGCC-3ʹ (reverse)

CAVIN1 Primers: 5ʹ-GGGCCGTAGACCAGATCCA-3ʹ (forward), 5ʹ-CTTGCTCACCGTATTGCTCGT-3ʹ (reverse)

CAVIN3 primers: 5’-CCGAAGAAGCTCTGCTCCAA-3’ (forward), 5’-GGGCGTGAGTGCTACATTCT-3’ (reverse)

CAV1 primers: 5’-TACGTAGACTCGGAGGGACAT-3’ (forward), 5’-GAAGCTGGCCTTCCAAATGC-3’ (reverse)

VEGFR2 primers: 5’-TGGTCAGGCAGCTCACAGTCC-3’ (forward), 5’-GTTCCGGTTCCCATCCTTCAATAC-3’ (reverse)

STARD4 Primer: 5ʹ-GATAAGTGGCGAGTTGCTAAGAA-3ʹ (forward), 5ʹ-ACACCTTGGGCTTTGTAGAGATA-3ʹ (reverse)

GAPDH primers (internal control): 5’-GGAGCGAGATCCCTCCAAAAT-3’ (forward), 5’-GGCTGTTGTCATACTTCTCATGG-3’ (reverse).

Reaction mastermix (8 μL) contained 5 μL supermix, 2.5 μL ddH2O (Bio-Rad), 0.5 μL primers (10 μM) was aliquoted into pre-cooled white PCR plates (Bio-Rad), followed by the addition of 2 μL of diluted cDNA (5 ng/μL) to each well, resulting in a total volume of 10 μL. The plate was loaded into the CFX thermal cycler with the program set for an initial activation of HotStart TAQ (Biolabs) at 95°C for 10 minutes, followed by 40 cycles of denaturation at 95°C for 15 seconds and annealing/extension at 60°C for 60 seconds. All reactions were performed in triplicate.

### Data plots

All visualizations were performed using custom scripts in R (v4.2.2) and Bioconductor (v3.18). The following R packages were used: gplots (v3.1.3) for heatmap.2(), ggplot2 (v3.4.2), ggbreak (v0.1.2), dplyr (v1.0.10), ggpubr (v0.6.0), enrichR (v3.2), clusterProfiler (v4.6.0), and Seurat (v4.3.0)

## Acknowledgements

MPC acknowledges support from the Oncode Institute, KWF (Dutch Cancer Society) Health∼Holland TKI Grant, NWO (the Netherlands Organization for Scientific Research) Veni and Vidi Grants, Stichting Ammodo Grant. We appreciate Josephine Nefkens Stichting’s support on the UFO microscope. RGR thanks the Ministry of Science, Research and Technology of the Islamic Republic of Iran, for the fellowship support. JFH thanks the Chinese Scholarship Council, for the fellowship support.

## Author contributions

RGR organized and conducted the HNSCC experiments, functional validation assays, immunostaining, and contributed to the identification of gene signature. JFH performed the functional validation assays and contributed to the discovery of protein trafficking and SRF’s role in HNSCC metastasis. MTLC scripted, performed and plotted the single-cell RNA sequencing analysis and contributed to the identification of gene signature. LY analyzed the migration data and helped with the BC-seq assay. JT and DB performed the single-cell RNA sequencing analysis and contributed to the identification of gene signature. TCC and LY performed the cell segmentation and tracking analyses. RGR, JFH and MTLC analyzed and plotted the migration inhibition, genetic interference and overexpression data. YB helped with the whole BC-seq pipeline and cell sorting. WB helped with semi-bulk BC-seq experiments. JaS contributed to the knockdown experiments. JAUH provided HNSCC samples and inputs on the experimental results. DV helped with the nCounter assays. TvdB and VvD helped with HNSCC FFPE sample preparation and annotation. RGR, JFH and MPC designed the experiments. RGR and MPC wrote the paper with input from all authors. MPC contributed to and supervised all aspects of the project.

