## Supplementary information for "Behavior-coupled single-cell transcriptomics reveals trafficking protein-dependent regulation of tumor cell migration"

\*These authors contributed equally

**Supplementary Figure S1-S13**

### Supplementary Figures

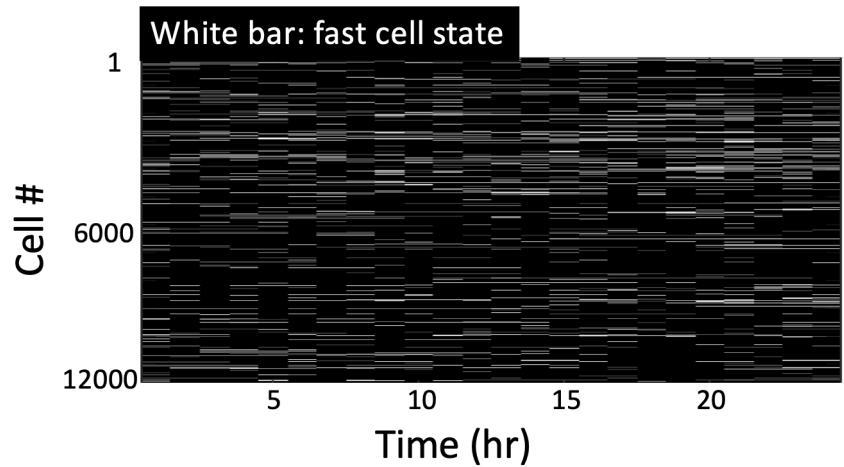

**Supplementary Figure S1. BC-seq identified FMCs.** The migratory state of 12,000 individual cells was tracked over the course of 24 hours. Cells in the fast migratory state are labelled in white, while those in the slow migratory state are labelled in black. Cells identified in the fast migratory state (called fast migrating cells (FMCs)) during a one-hour time-lapse recording were subsequently isolated (within 30 minutes after identification) and subjected to single-cell RNA sequencing using the BC-seq pipeline.

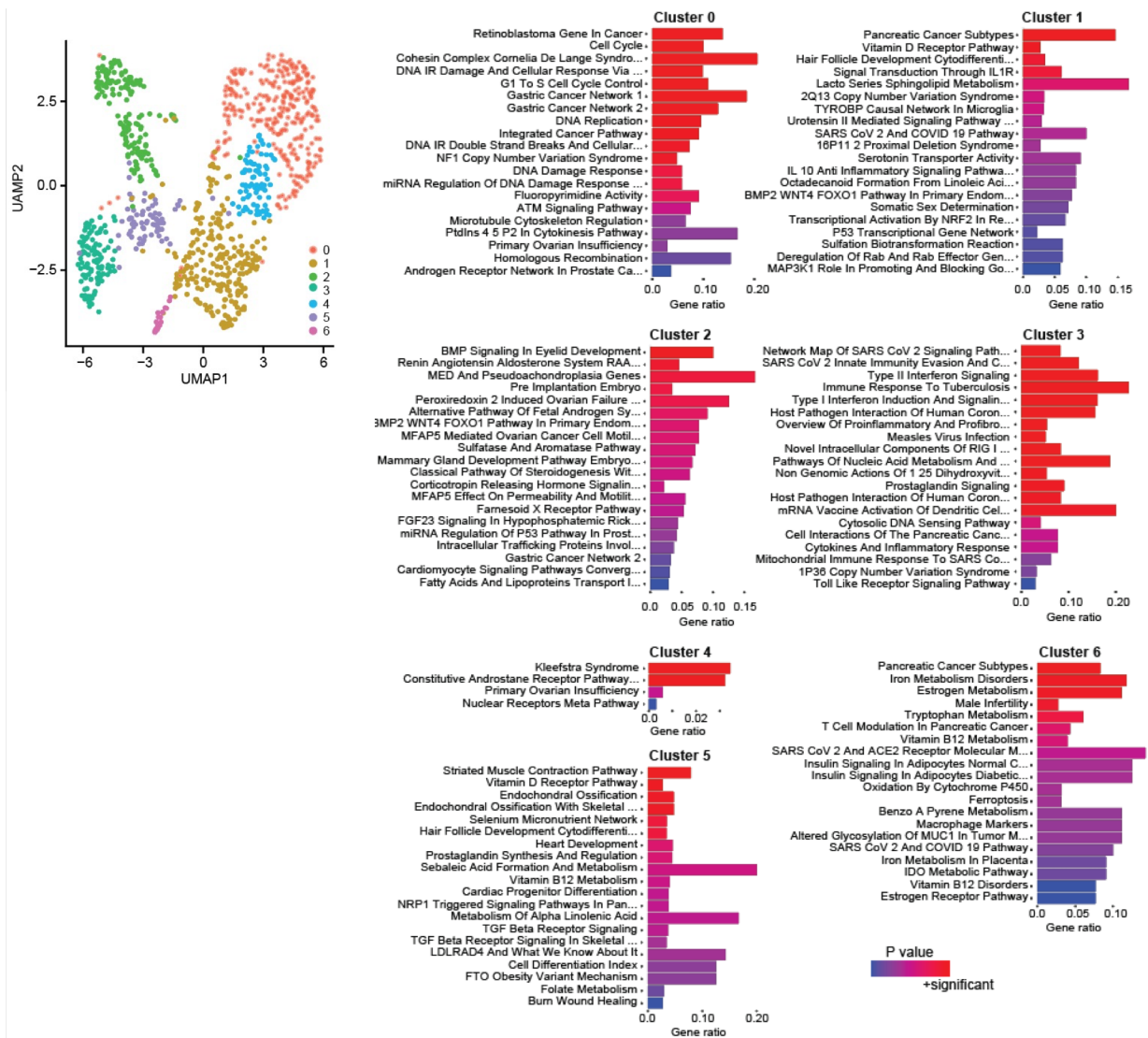

**Supplementary Figure S2. WikiPathway analysis of the seven clusters (Cluster 0-6) identified through Seurat-based clustering analysis (non-BC-seq scRNAseq data). Fisher's exact test was applied for the Enrichr pathway enrichment analysis.**

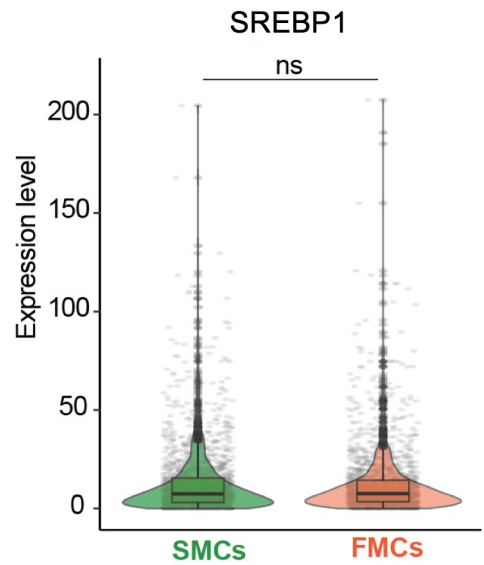

**Supplementary Figure S3. A violin plot shows gene expression level of SREBP1 for SMCs and FMCs. *P* values were calculated using the Wilcoxon rank sum test with Bonferroni correction.**

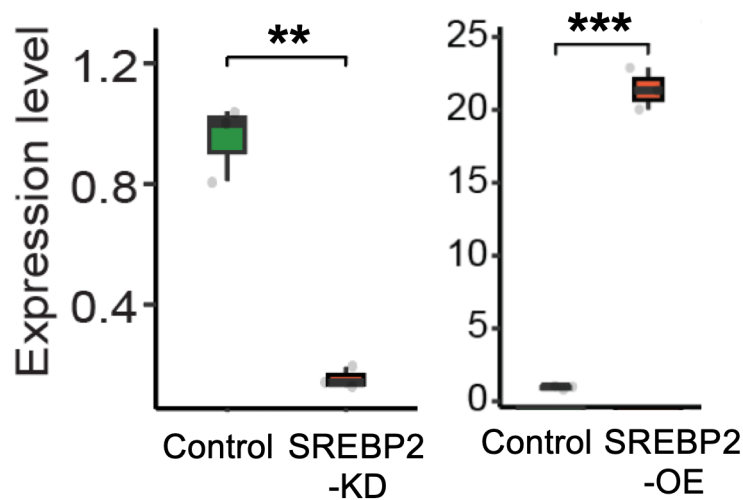

**Supplementary Figure S4. qPCR results for the SREBP2 gene under the SREBP2-knockdown (KD) and -overexpression (OE) conditions, compared to the control. Expression levels were normalized to GAPDH (housekeeping gene), and *P* values were calculated using a two-tailed unpaired *t*-test.**

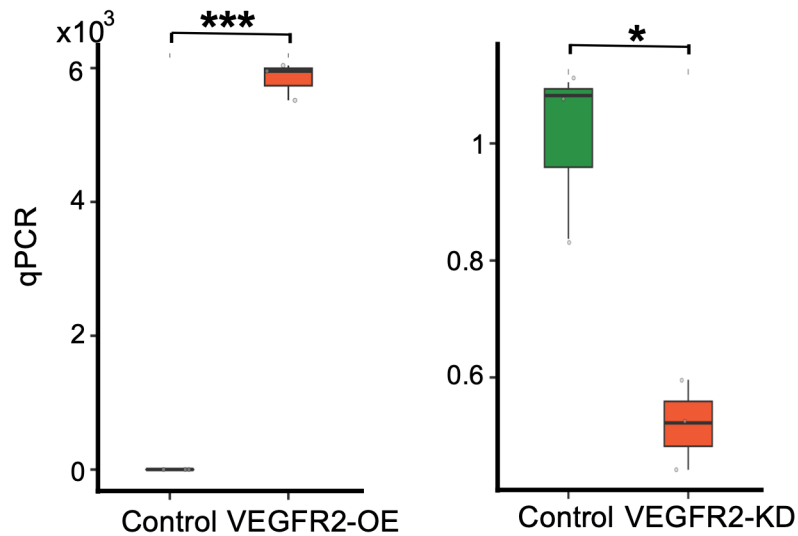

**Supplementary Figure S5. qPCR results for the VEGFR2 gene under the VEGFR2-overexpression (OE) and VEGFR2-knockdown (KD) conditions compared to the control.** Expression levels were normalized to GAPDH (housekeeping gene), and *P* values were calculated using a two-tailed unpaired t-test.

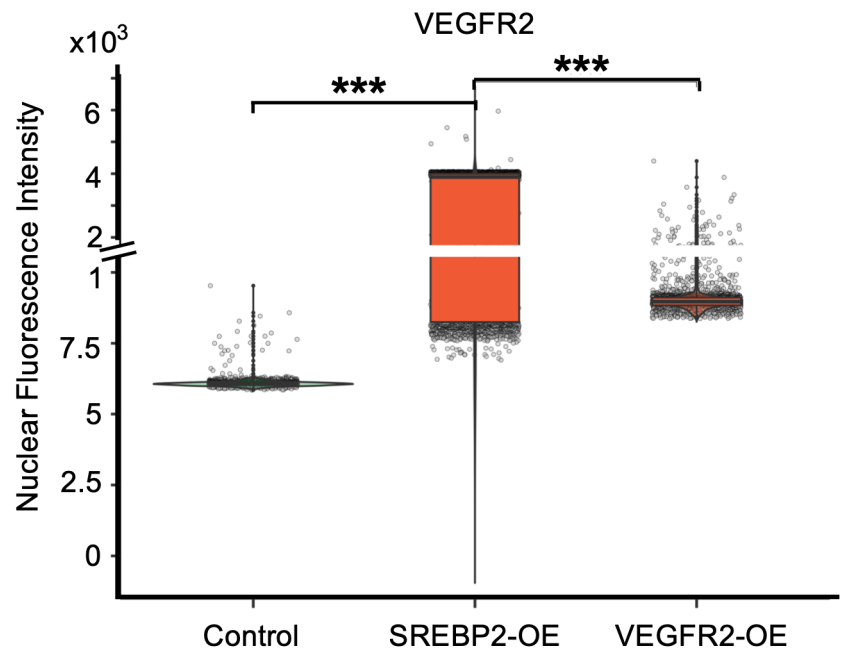

**Supplementary Figure S6. Nuclear immunofluorescence intensity of VEGFR2 under control, SBREP2-overexpression (OE), and VEGFR2-OE conditions.** *P* values were calculated using the Wilcoxon rank sum test with Bonferroni correction.

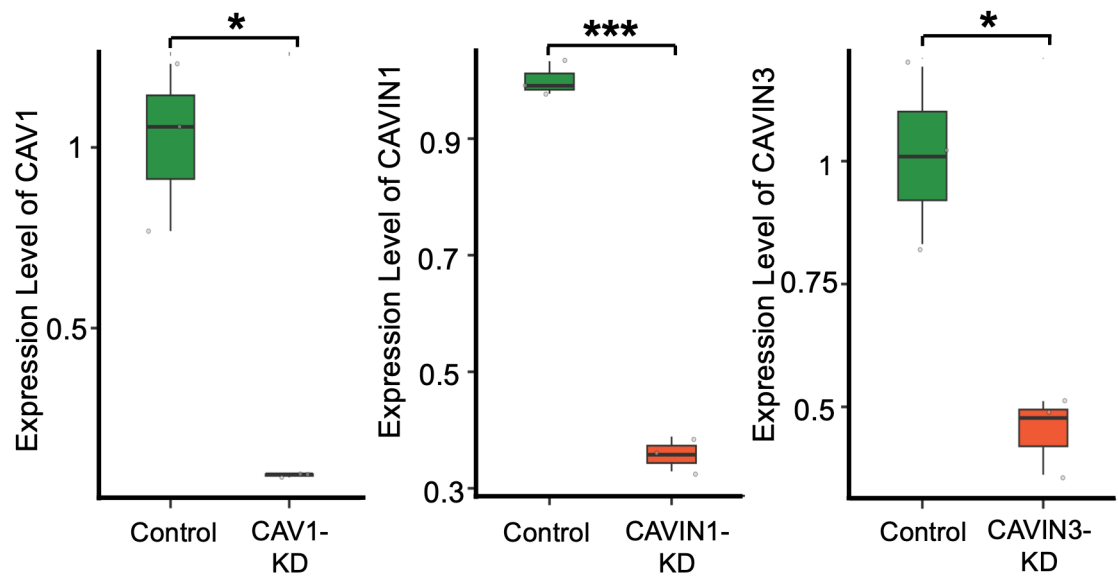

**Supplementary Figure S7.** The qPCR quantification of CAV1, CAVIN1 and CAVIN3 gene expression following gene knockdown (KD). *P* values were calculated using a two-tailed unpaired t-test.

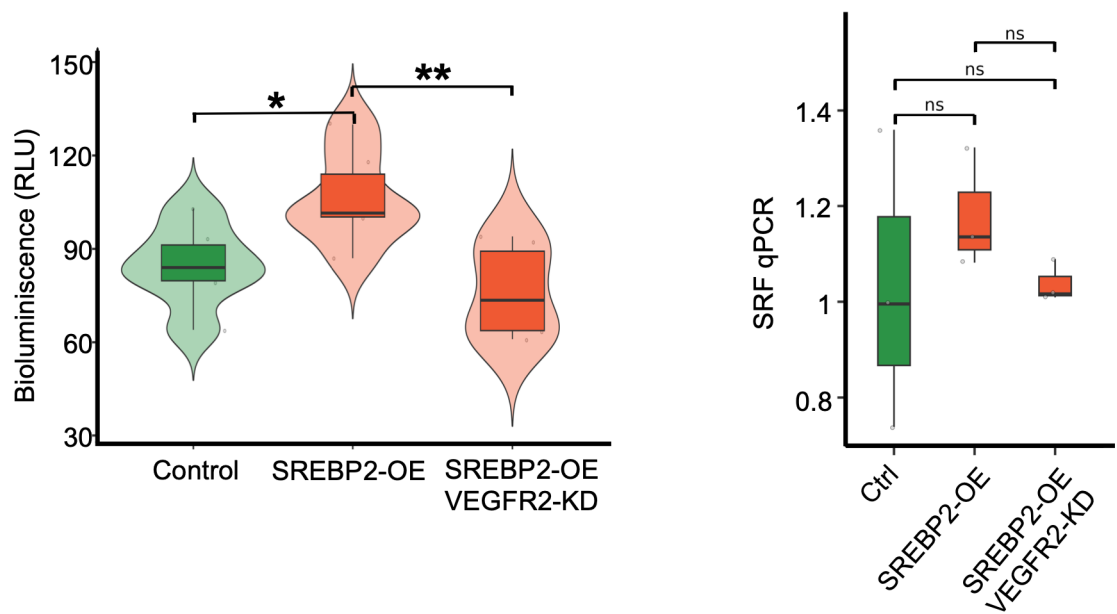

**Supplementary Figure S8.** (Left panel) Bioluminescence intensity (relative light units, RLU) of cells under control, SREBP2-overexpression (OE), and combined SREBP2-OE with VEGFR2-knockdown (KD) conditions. *P* values were calculated using the Wilcoxon rank sum test with Bonferroni correction. (Right panel) The qPCR quantification of serum response factor (SRF) under control (Ctrl), SREBP2-OE, and combined SREBP2-OE with VEGFR2-KD conditions. *P* values were calculated using a two-tailed unpaired t-test.

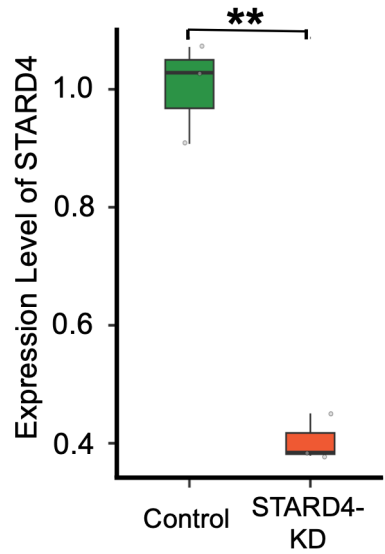

**Supplementary Figure S9. The qPCR quantification of STARD4 gene expression following gene knockdown (KD).** *P* values were calculated using a two-tailed unpaired t-test.

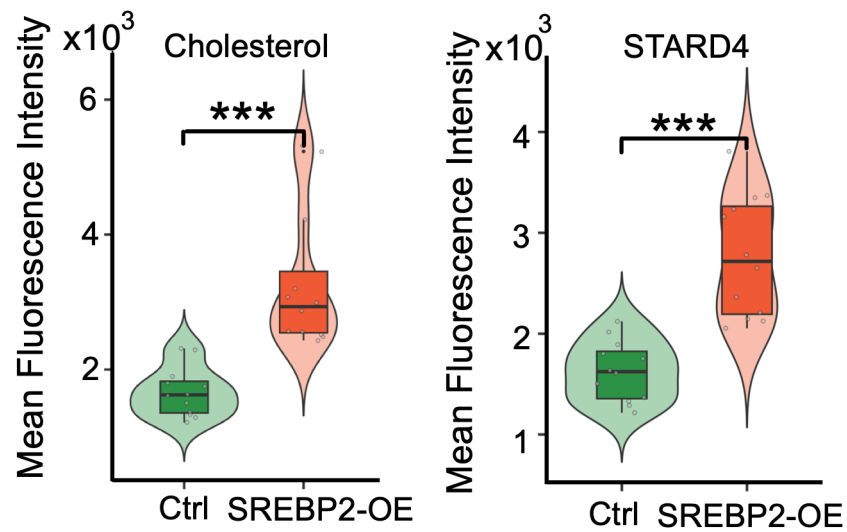

**Supplementary Figure S10. Fluorescence intensity of cholesterol (left panel) or STARD4 (right panel) signals under control (Ctrl) and SREBP2-overexpression (OE) conditions.** *P* values were calculated using the Wilcoxon rank sum test with Bonferroni correction.

|  |  |  |  |
| --- | --- | --- | --- |
| ACAT2 | DYRK3 | MSMO1 | RRAS |
| ACTA2 | EDEM1 | MSN | RSU1 |
| ALDH1A3 | FBLIM1 | MYH9 | SEC24D |
| ACLY | FDFT1 | MYL12A | SERPINE2 |
| ACTN1 | FDPS | NSDHL | SFN |
| ACTR3 | FN1 | OCIAD2 | SMAD3 |
| ALOX15B | GPAT3 | PALM2AKAP2 | SNAI2 |
| ANXA5 | HAPSTR1 | PAK2 | SNX9 |
| ARHGEF2 | HMGCS1 | PDLIM7 | SNX12 |
| ARPC5 | HSPA1A | PFN1 | SOX15 |
| ARPC1A | HINT1 | PJA2 | SPARC |
| ATXN2 | IL1A | PROCR | SQLE |
| AQP3 | INSIG1 | PRRC1 | STARD4 |
| BZW1 | IRX2 | PNPLA8 | TACSTD2 |
| CAVIN1 | ID1 | PPP1R18 | TFE3 |
| CAVIN3 | ITGA5 | PPP2R1A | TGFB1I1 |
| CAP1 | LASP1 | PRSS23 | TIMP1 |
| CCNG2 | LGALS1 | PTGS2 | TJP3 |
| CD99 | LIMA1 | PTHLH | TPM1 |
| CDKN2A | LPIN1 | PTK2B | TPM4 |
| CERS2 | MAML2 | PXN | TUBB6 |
| COL1A1 | MAPKAPK2 | RAC2 | TUBB2A |
| COL1A2 | MYO5A | RBM3 | VASP |
| DDIT3 | MBOAT2 | RHOC | WNT7A |
| DNAJB9 | MDK | RNF11 | WSB1 |

**Supplementary Figure S11.** A total of 100 gene markers were assayed using the nCounter platform. 39 significant genes were identified via univariate tests (highlighted in red and green). A 15-gene signature (highlighted in red) was identified through 50 iterations of bootstrap resampling with replacement, using a logistic regression model with Elastic Net regularization.

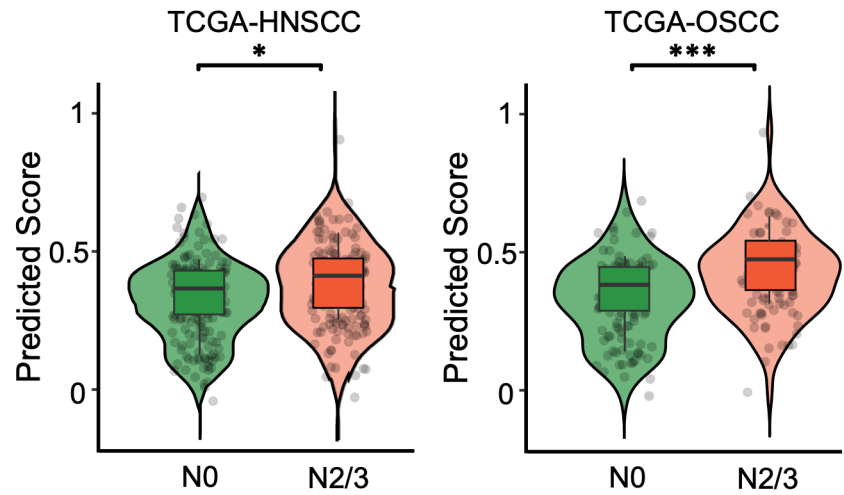

**Supplementary Figure S12. Predicted scores using the 15-gene signature on patients from the TCGA database.** (Left) Predicted scores of N0 and N2/3 HNSCC patient samples from TCGA calculated using a logistic regression model with Elastic Net regularization. (Right) Predicted scores of N0 and N2/3 OSCC patient samples from TCGA calculated using a logistic regression model with Elastic Net regularization. P values were calculated using the Wilcoxon rank sum test with Bonferroni correction.

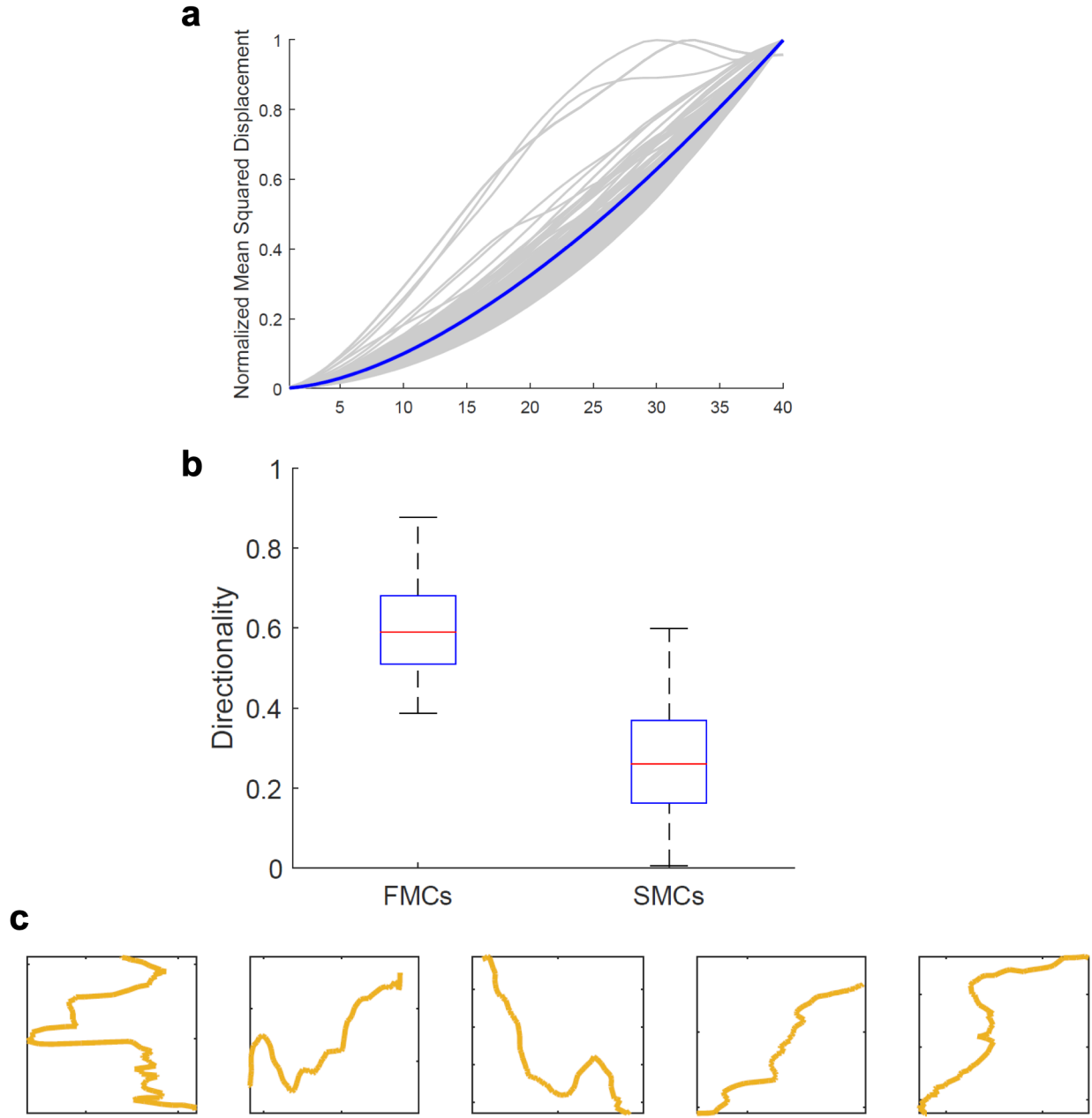

**Supplementary Figure S13. Characterizations of FMCs.** **a.** Normalized mean squared displacement (MSD) of FMCs. MSD per cell is shown in gray, and the mean trend is shown in blue. **b.** Directionality of FMCs and SMCs. Directionality measures the deviation between the cumulative distance $S_{total}$ (i.e., total distance travelled via all time-points) and linear distance $S_{linear}$ from the start to end time-point. The ratio is given as $r=S_{linear}/S_{total}$. **c.** Single cell migratory trajectories of five representative FMCs.
